# A Developmental Single-Cell Atlas of the *Drosophila* Visual System Glia Reveals Cell Type Diversification and Subcellular mRNA Compartmentalization

**DOI:** 10.1101/2023.08.06.552169

**Authors:** Amanda A. G. Ferreira, Sergio Córdoba, Raghuvanshi Rajesh, Ben Jiwon Choi, Claude Desplan

## Abstract

Glial cells are essential for proper nervous system development and function. To understand glial development and function, we comprehensively annotated the glial cells from two large single-cell mRNA-sequencing (scRNA-seq) atlases of the developing *Drosophila* visual system. This allowed us to identify all glial cell types from larval to adult stages and follow their developmental trajectories to understand how the diversity of glial types is generated during development. We show that whereas most glial types, such as chiasm glia, gradually change their transcriptome as they mature during development, neuropil glia that transcriptionally appears as a single cell class in larvae, splits into ensheathing (EG) and astrocyte-like (ALG) glia types during pupal stages. We have experimentally validated these developmental trajectories and identified the genetic markers expressed through the differentiation between EG and ALG classes. Unexpectedly, our analysis of scRNA-seq datasets allowed us to discover that the transcriptome of glial cell bodies can be distinguished from that of their processes. We have identified that processes are enriched for distinct mRNAs that were validated in vivo. This work provides the most detailed transcriptomic analysis of optic lobe glia during development and helps explain the expansion of glial diversity observed in the adult visual system. We also present an innovative computational approach to identify mRNA species that are differentially localized to cell bodies or cellular processes.

## Introduction

Glial cells play a fundamental role in nervous system development and function. These cells coordinate neural stem cell division, neuronal differentiation, axon pathfinding, and synapse formation^1–4^. Glial cells also serve as structural and metabolic support, clean apoptotic cells and cell debris, insulate axons, actively participate in synapses, and constitute the blood-brain barrier^5–8^. Dysfunctional glia is associated with several diseases, such as multiple sclerosis, glioblastoma, Alzheimer’s disease, Parkinson’s disease, autism, and psychiatric disorders^4,9,10^.

Anatomically, glia are highly polarized cells that have extensive cellular processes in which mRNAs are actively localized in specific subcellular compartments^11–13^. For example, the mRNAs encoding the major myelin protein MBP, are found concentrated in the processes of glial cells surrounding the axon they ensheath, and their translation is triggered by the activity of the neuron^14^.

The *Drosophila* optic lobe has four main classes of glia: surface (perineurial & subperineurial), cortex, neuropil, and nerve-wrapping glia^8,15–17^. Surface glia surrounds the brain and form the blood-brain barrier that controls exchanges between the brain and the hemolymph^18^. Cortex glia is in close contact with the neuronal cell bodies and its function is to nourish neurons and eliminate dead ones ^7^. Neuropil glia is associated with neuronal terminals and is essential for synapse formation and neurotransmitter recycling^19,20^. Finally, nerve-wrapping glia wrap around axon bundles that cross from one neuropil to another, serving as support and protection for these neurons^15,17,21^. Glial cells are also involved in the development of the nervous system, contributing to axon pathfinding and neuronal circuit assembly^22^. Cortex glia influence the proliferation of neural stem cells (neuroblasts in flies) and the neuroepithelium to neuroblast transition^1–3^. Previous efforts to characterize glial cells in the *Drosophila* brain have relied on enhancer traps to identify different glial types based on their morphology^15,23^. More recently, the transcriptome of the glial cells of the adult optic lobe has been annotated^17^. However, the glial diversity in the larval brain and how it evolves through development until adulthood has not yet been investigated.

To comprehensively characterize the transcriptomic diversity of glia in the fly optic lobe and study glial development from larval to adult stages, we leveraged different scRNA-seq datasets at different stages of development that were previously generated in the laboratory^24,25^. About 32,000 sequenced optic lobe glial cells from larvae, four pupal stages, and adults were analyzed. We annotated most glial cells and validated these annotations at the third instar larval (L3) and adult stages. Their cell identities were then transferred to all pupal stages by integrating the datasets^24^. Unexpectedly, we have discovered that the mRNA expression levels of the classical glial marker *repo* are generally low and its expression alone is insufficient to identify all glial clusters in scRNA-seq datasets. We identified three new genes with broad expression among glial cell types (*CG32032*, *AnxB9*, and *GstE12)* that can be used along with *repo* to unambiguously identify all glial clusters. We also found that glial types specified in the larva follow three differentiation trajectories through adulthood: (1) glial types whose transcriptome minimally changed from larvae to adults, such as cortex and perineurial glia; (2) glial types whose transcriptome gradually changed and formed a linear trajectory, like inner and outer chiasm glia; and (3) glial types that formed an initial linear trajectory and then split into at least two different fates, such as medulla neuropil glia, which differentiates into astrocyte-like/ensheathing glia, and lamina neuropil glia that gives rise to epithelial/marginal glia in the adult. Moreover, while analyzing these scRNA-seq datasets, we discovered that certain glial cells were broken down during cell dissociation into different cell fragments that were then sequenced independently and clustered separately: Unique glial cell types are thus represented by multiple clusters corresponding to the cell bodies and cell processes with distinct mRNA contents.

## Results

### Identification of new glia-specific markers

Marker genes are essential to identify glial clusters in scRNA-seq datasets. In *Drosophila*, glial cells are most often defined *in vivo* by the expression of the Repo protein^26^. To identify glial clusters in scRNA-seq, we selected the larval clusters^25^ whose transcriptome exhibited a high correlation with the transcriptome of fluorescence-activated (FAC)-sorted cells based on the expression of a *repo*-GFP transcriptional reporter, and a low correlation with that of neurons FAC-sorted based on *Elav* expression, a pan-neuronal marker, as previously done for the adult optic lobe dataset^24^ (**Fig. 1A** and **S1A**). Surprisingly, the endogenous *repo* mRNA was lowly expressed and often absent in many of the glial clusters: *repo* mRNA was only significantly expressed in 11 out of 23 larval glial clusters and in 5 out of 21 adult glial clusters (**Fig. 1B, C**). This low frequency of transcript detection in scRNA-seq experiments is probably due to ‘dropouts’ that are common for lowly expressed genes^27,28^. Consequently, the absence of *repo* detection in glial clusters in scRNA-seq data may result in many clusters mistakenly not being considered glial cells. For example, in the larva both types of surface glia (subperineurial, spg and perineurial, pg) had low *repo* expression levels in the dataset (cluster 21 for spg, and 8 and 14 for pg, **Fig 1B**), which correlates to the low detection of *repo* mRNA by Hybridization Chain Reaction RNA Fluorescence *in situ* Hybridization (HCR RNA FISH)^29,30^ even if these cells could easily be identified as glial cells by antibody detection of the Repo protein (**Fig. S1D**).

**Figure 1.**
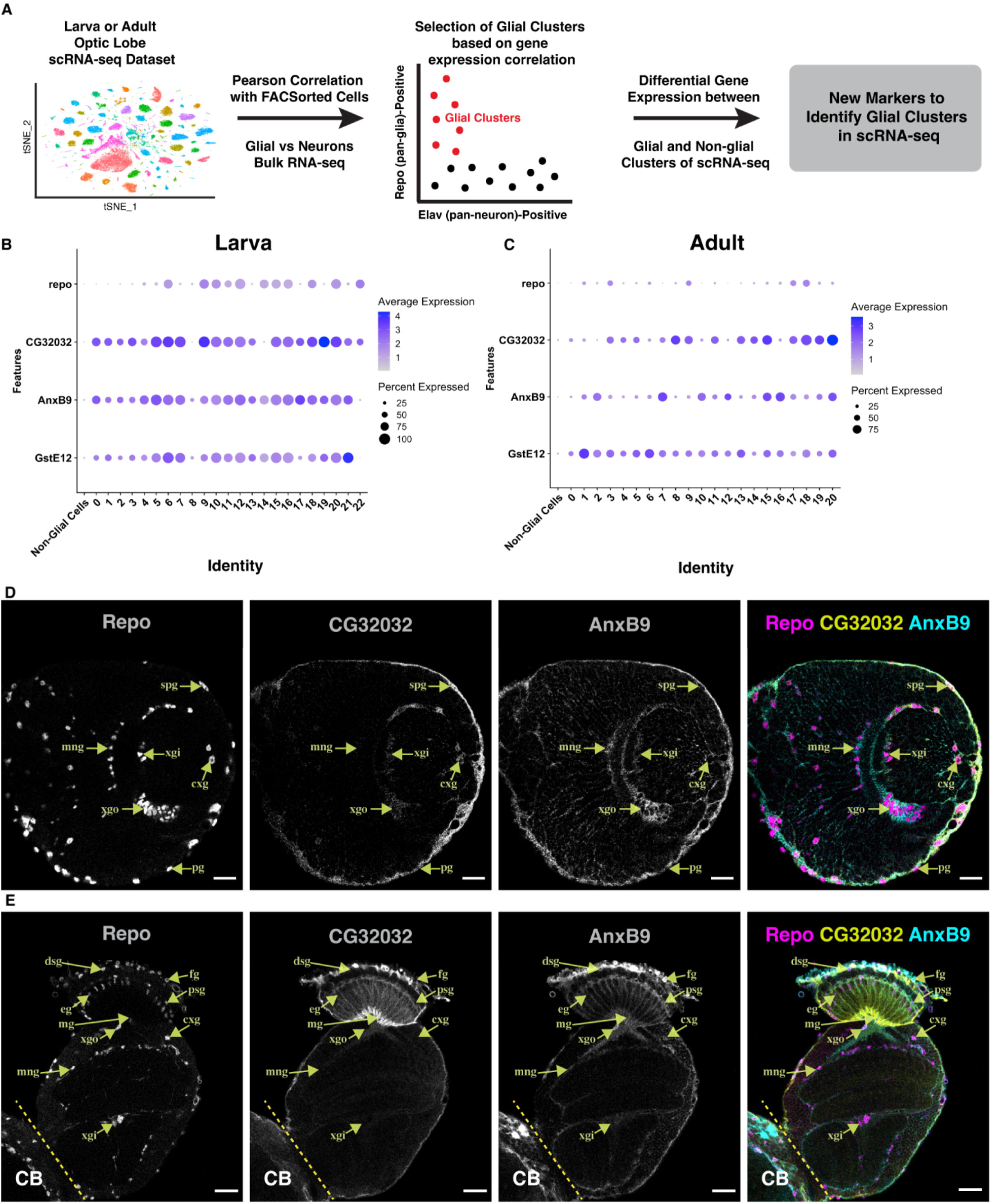
Identification of New Glia-Specific Marker Genes. **(A)** Schematic representing how glial cell clusters were defined. Pearson correlation between the transcriptome of the clusters with that of bulk transcriptome of *repo*-positive and *elav*-positive cells was used to define glial clusters. The defined clusters were used to find new glial markers. **(B)** Dot plot showing the newly identified glial markers in the larva (L3). Numbers identify individual glial clusters, and “Non-Glial Cells” represent the combined gene expression of all the remaining cells from the whole larva dataset from Konstantinides *et al.*, 2022. The color of the dots indicates the mean normalized gene expression across all cells in a group (Average Expression), while the dot size indicates the fraction of cells with non-zero expression (Percent Expressed). **(C)** Dot plot showing the newly identified glial markers in the adult dataset per glial cluster. “Non-Glial Cells” represent the combined gene expression of all the remaining cells from the whole adult stage dataset from Özel *et al.*, 2021. **(D)** Immunostaining of the larval (L3) optic lobe showing the localization of Repo, CG32032 and AnxB9 antibodies. **(E)** Immunostaining of the adult optic lobe showing the localization of Repo, CG32032 and AnxB9 antibodies. **CB** = Central Brain, **fg** = fenestrated glia, **pg** = perineurial glia, **spg** = subperineurial glia, **dsg** = distal satellite glia, **psg** = proximal satellite glia, **eg** = epithelial glia, **mg** = marginal glia, **xgo** = outer chiasm glia, **xgi** = inner chiasm glia, **cxg** = cortex glia, **mng** = medulla neuropil glia, **rwg** = retina wrapping glia. Scale bar = 20 µm.

To overcome this challenge, we looked for other glia-specific genes with higher expression that could allow us to robustly identify glia in scRNA-seq datasets, independently or together with *repo* expression. By comparing all the larval glial clusters with the other non-glial clusters, we identified several genes that were differentially expressed in glial cells as compared to neurons: *Ama*, *MtnA*, *AnxB9*, *CG32032*, *CG7433*, *GstE12*, *CG9686*, *CG15209*, and *Npc2a* (**Fig. S1B, C**). Among these genes, *AnxB9*, *CG32032* and *GstE12* were individually expressed in most glial clusters, including those that did not express *repo* (**Fig. 1B, C** and **Fig. S1C**) but were largely absent in neuronal clusters, with few exceptions. *AnxB9* belongs to the family of annexins that bind to phospholipids and participate in membrane remodeling, membrane trafficking and exocytosis^31^. Annexins are widely found in the nervous system and show different expression patterns in neurons and glia^32,33^*. CG32032* is predicted to belong to the calycin protein superfamily (FlyBase^34^), which includes lipocalins and fatty acid-binding proteins often found in glial cells, such as the human *FABP7* and *PMP2*^6,35^. Finally, *GstE12* encodes a cytoplasmic Gluthatione S-Transferase (GST) enzyme^36^. In mice, astrocytes enhance proinflammatory response in the brain through GSTs, and the knockdown of these enzymes in astrocytes compromises microglial activation^7^.

We generated antibodies for AnxB9 and CG32032, both of which showed highly glia-specific signal in immunostainings, which confirmed that the protein product of *CG32032* and *AnxB9* genes is present in glial cells in both L3 larvae and adults (**Fig. 1D, E**). However, we observed some cell-type specific variability: in L3 brains AnxB9 staining was high in giant chiasma glia (xgo) and low in epithelial and marginal glia (eg/mg), while CG32032 antibody signal was low in xgo and high in eg/mg (**Fig. S1E, F**). Thus, *CG32032* and *AnxB9* are both expressed strongly and mostly specifically in glia and, in combination with *repo*, are excellent candidates for the identification of glial cell clusters in scRNA-seq datasets. Based on the expression of at least two of these genes, all previously identified glial clusters in both larval (23 out of 23 clusters) and adult (21 out of 21 clusters) datasets were identified, and none of the non-glial clusters expressed these combinations. To test whether these genes could help identify glial types in other datasets, we analyzed their expression in a published optic lobe scRNA-seq dataset from pharate adults (96 hours pupation – P96)^37^, and found that *repo* was lowly expressed in three clusters annotated as glia by the authors (**Fig. S1G**). However, these three clusters expressed at least two of the new glial markers. Moreover, this dataset contained clusters called “C clusters” that were annotated neither as glia nor as neurons. Many of the cells in these “C clusters” expressed the new glial markers. Indeed, out of all 321 cells in the “C clusters”, 103 had transcriptomes that resembled adult subperineurial glial cells. Similarly, cluster N183, previously annotated as an unknown neuronal cluster, also showed expression of these glial markers and was composed of 19 cells, 9 of which also correlated with subperineurial glial cells (**Fig. S1G**). Another published *Drosophila* scRNA-seq dataset of the adult ventral nerve cord (VNC) was also analyzed^38^. Two of the clusters that the authors had identified as glia/neuron mix due to low expression of *repo* could be confidently identified as glia when using the combined expression of *repo*, *CG32032*, *AnxB9,* and *GstE12* (**Fig. S1H**, arrows) and indeed the authors later managed to identify these clusters as glia, based on specific glial subtypes markers^39^. Interestingly, in this VNC dataset a number of clusters that did not express any of the other glial markers did exhibit *AnxB9* expression. Accordingly, we observed strong AnxB9 antibody signal in non-glial cells, presumably neurons, in the larval VNC (**Fig. S1I**). In summary, using four different datasets, we showed that *repo*, *CG32032*, *AnxB9,* and *GstE12* are marker genes that can be used in combination to consistently identify glial clusters in *Drosophila* scRNA-seq datasets.

### Annotation of glial clusters in a developmental scRNA-seq atlas of the fly optic lobe

Next, we sought to evaluate glial diversity in the larval and adult stages. High-resolution transcriptomic atlases of the optic lobe throughout development^24,25^ had identified more than 200 cell types, including progenitors, neurons and glia, but glial clusters were not annotated as specific glial types. We excluded neurons and re-clustered the *bona-fide* glial cells (**Fig. 2B**)^40^. Glial cell bodies occupy stereotypical positions in the brain and consistently express Repo protein, which allowed us to match specific gene expression from each cluster to known glial cell types. We annotated and validated larval clusters based on known^7,15,20,23,39,41–48^ or new markers identified in this study (summarized in **Table 1** and **Methods**) using transcriptional reporters, protein traps, enhancer traps, antibodies, as well as specific HCR RNA FISH probes (**Fig. 2A** and **C-M**).

**Figure 2.**
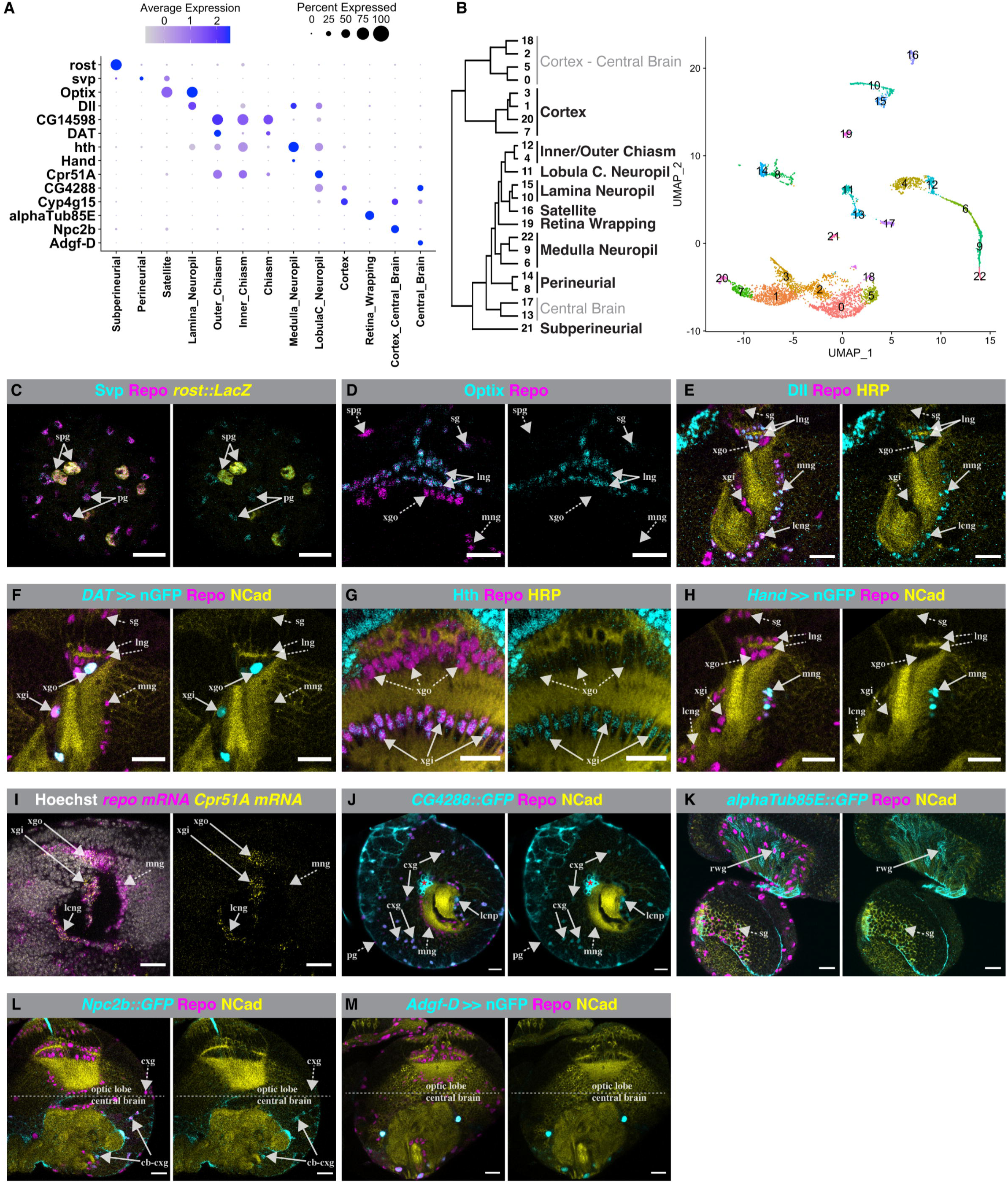
Cluster Validations for Larva Glial Annotations. **(A)** Dot plot showing the expression levels of the genes used for the validation of each annotated cluster in L3. **(B)** Left, hierarchical tree showing cluster annotations after validations reveals 10 cell types (in bold text), excluding cells of central brain origin. Note that Chiasm glia was manually separated into Inner and Outer Chiasm subtypes. Right, unsupervised clustering of larval glial cells. **(C-M)** Immunostaining and FISH analysis of marker gene expression in the L3 optic lobe. Neuropil structure is revealed by N-cadherin (Ncad) or HRP staining (yellow), while Repo (magenta) is used to label glia nuclei in the left panels. Marker gene expression is shown using antibodies, *lacZ* reporters, protein tags or specific Gal4 lines driving UAS-nGFP expression (cyan). **(C)** *rost-lacZ* reporter (yellow) is only expressed in subperineurial glia. Anti-Svp antibody (cyan) labels both perineurial and subperineurial glia. **(D)** Optix (cyan) is present in satellite, epithelial, and marginal glia but not in subperineurial, outer chiasm and medulla neuropil glia. **(E)** Dll (cyan) labels epithelial, marginal, lobula complex neuropil and medulla neuropil glia but not satellite, outer, and inner chiasm glia. **(F)** *DAT*-Gal4 (cyan) is expressed in the outer and inner chiasm glia. It is less frequently observed in the inner chiasm glia. **(G)** Hth (cyan) antibody is higher in inner chiasm than in outer chiasm glia. **(H)** *Hand*-Gal4 (cyan) is only expressed in medulla neuropil glia. **(I)** HCR RNA FISH probes against *Cpr51A* (yellow) and *repo* (magenta) mRNA. Hoechst antibody is shown in grey. *Cpr51A* is expressed in the chiasm and lobula complex glia. The image shows the standard deviation projection of three consecutive focal planes 2 µm apart. **(J)** GFP-tagged *CG4288* (cyan) is expressed only in the cortex and lobula complex glia but not in the perineurial, medulla neuropil and inner chiasm glia. Asterisk indicates Bolwig organ. **(K)** GFP-tagged *alphaTub85E* (cyan) is expressed in retina wrapping glia. **(L)** GFP-tagged Npc2b (cyan) is expressed only in cortex glia from the central brain. **(M)** Adgf-D (cyan) is expressed in the central brain glia. **pg** = perineurial glia, **spg** = subperineurial glia, **sg** = satellite glia, **eg/mg** = epithelial and marginal glia, **mng** = medulla neuropil glia, **xgo** = outer chiasm glia, **xgi** = inner chiasm glia, **cxg** = cortex glia, **rwg** = retina wrapping glia, **cb-cxg** = central brain cortex glia, **lcng** = lobula complex neuropil glia. Gal4-UAS (*>>*), *GFP/LacZ* from construct (::). Scale bar = 20 µm.

At the adult stage, we based our annotations on cell-type specific markers established by earlier studies that had identified genes expressed in specific subtypes of glia using enhancer traps, antibody staining, cell-type specific RNA-seq, and scRNA-seq ^20,39,46,49–54^. We determined the identity of adult clusters by taking advantage of the spatial distribution of glial cells combined with immunohistochemistry for genes that were expressed in one or a few clusters (**Fig. 3B** and **C-L**; see **Table 2** for the list of markers, annotations, and corresponding references). Ultimately, we could assess the transcriptome of all glial cell types and determine specific markers for each of them (**Table S1**). This represents a powerful resource for studying glial cells and their contribution to the development and function of the optic lobe.

**Figure 3.**
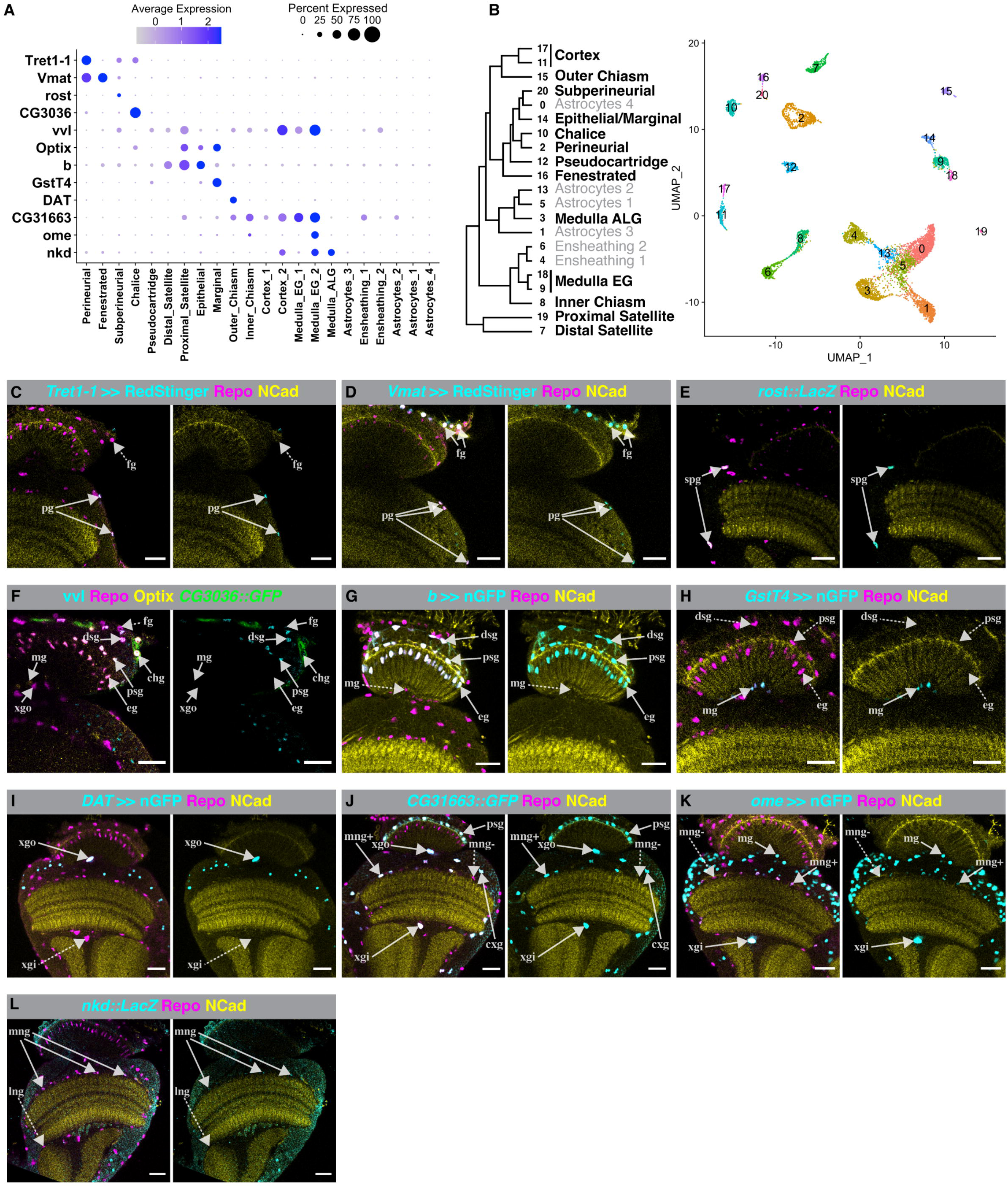
Cluster Validations for Adult Glial Annotations. **(A)** Dot plot showing the expression levels of the genes used for the validation of each annotated cluster in adult. **(B)** Left, hierarchical tree showing cluster annotations after validations reveals 14 distinct cell types (in bold text) plus a number of clusters annotated preliminarily as Astrocyte- and Ensheathing-like glia. Right, unsupervised clustering of adult glial cells. **(C-L)** Immunostaining analysis of adult stage optic lobe. Neuropil structure is revealed by Ncad or HRP staining (yellow), and Repo (magenta) labels glia nuclei in the left panels. Marker gene expression is shown using antibodies, *lacZ* reporters, protein tags or specific Gal4 lines (cyan). **(C)** *Tret1-1*-Gal4 (cyan) is only expressed in perineurial glia and not in fenestrated. **(D)** *Vmat-*Gal4 (cyan) is expressed in both perineurial and fenestrated glia. **(E)** *rost*-*lacZ* reporter (cyan) is expressed only in subperineurial glia. **(F)** GFP-tagged *CG3036* (green) is expressed only in chalice glia. Vvl antibody (cyan) labels distal and proximal satellite glia. Optix (yellow) is present in chalice, marginal, epithelial and proximal satellite glia. **(G)** *b*-Gal4 (cyan) is expressed in distal and proximal satellite glia and in epithelial glia, but not in marginal. **(H)** *GstT4*-Gal4 (cyan) is expressed only in marginal glia. **(I)** *DAT*-Gal4 (cyan) is expressed in outer chiasm glia but only seen rarely in inner chiasm glia. **(J)** GFP-tagged *CG31663* (cyan) is expressed in proximal satellite glia and some medulla neuropil glia. This might reflect ensheathing- (mng+) *vs.* astrocyte-like (mng-) identity. It is also expressed in outer and inner chiasm glia and in cortex glia. **(K)** *ome*-Gal4 (cyan) is expressed in inner chiasm glia, marginal glia and some medulla neuropil glia, which could reflect ensheathing (mng+) vs astrocytes-like (mng-) identity. **(L)** *nkd*-*lacZ* reporter (cyan) is expressed in medulla neuropil glia but not in lobula neuropil glia. **fg** = fenestrated glia, **pg** = perineurial glia, **spg** = subperineurial glia, **dsg** = distal satellite glia, **psg** = proximal satellite glia, **chg** = chalice glia, **eg** = epithelial glia, **mg** = marginal glia, **xgo** = outer chiasm glia, **cxg** = cortex glia, **lng** = lobula neuropil glia, **mng** = medulla neuropil glia. Gal4-UAS (*>>*), *GFP/LacZ* from construct (::). Scale bar = 20 µm.

### Developmental trajectories of glial cells from larva to adult stages

In the course of annotating glial types in larva and adult stages, we noticed that the genetic markers of glial types changed over time, and that there was a significant increase in glial cell diversity in the optic lobe, from 10 cell types in larva to at least 14 distinct cell types in adult (**Fig. 2B** and **Fig. 3B**). To investigate this increase, we integrated all glial cells from larvae^25^, P15 (15 hours after pupation), P30, P50, P70, and adults ^24^ into one dataset (**Fig. 4A**). Interestingly, the same cell types across stages clustered together, such as cortex, perineurial, subperineurial, lamina neuropil, medulla neuropil, inner and outer chiasm glia (**Fig. S2A, B**). The transcriptomic similarity of the same cell types throughout development corroborates the robustness of our marker-based cell type annotation (**Tables 1** and **2**).

**Figure 4.**
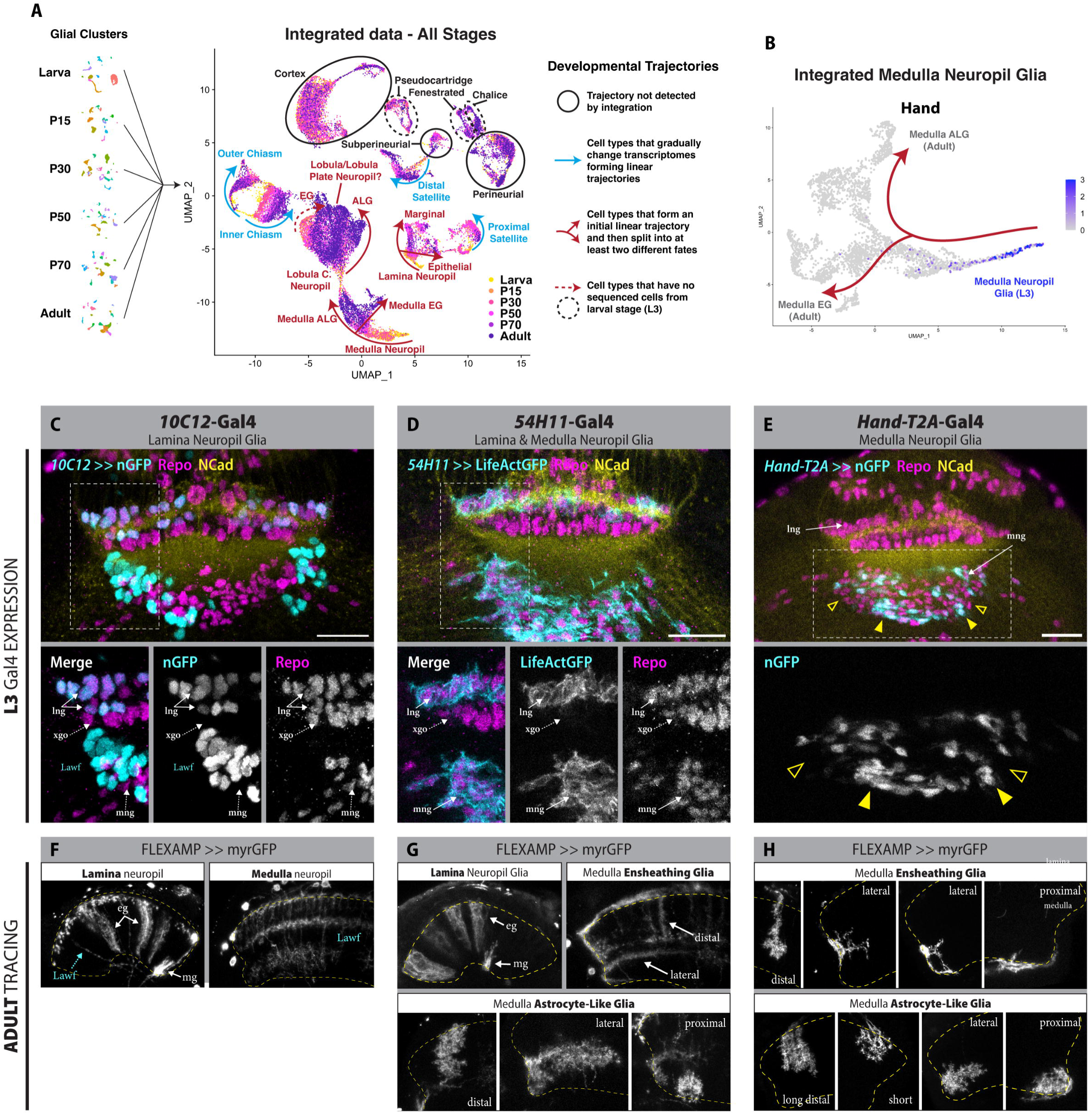
Developmental Diversification of Neuropil Glia. **(A)** Left, UMAPs representing all glial clusters from L3, P15, P30, P50, P70 and adult stage optic lobes. Right, UMAP integration of all developmental stages; cells are color-coded according to their stage of origin. Blue arrows highlight glial cells that show a gradual developmental trajectory from L3 to adult in our integration. Red arrows indicate trajectories where glial cells that are initially similar bifurcate into different terminal fates. Circled in black are glial cells that do not show a developmental trajectory from L3 to adult according to the integration. Dashed lines represent trajectories that do not contain cells from larval stage. **(B)** UMAP showing isolated medulla neuropil glia from the integrated dataset in **(A)**, where the divergent trajectories of medulla astrocyte-like (ALG) and ensheathing glia (EG) can be observed. *Hand* displays a salt-and-pepper expression that is restricted at the beginning of the trajectory (L3). **(C-E)** L3 expression of *10C12*-Gal4, *54H11*-Gal5 and *Hand-T2A*-Gal4 lines, driving the UAS-nlsGFP-PEST **(C** and **E)** or UAS-LifeActGFP **(D)** in cyan. Ncad (yellow) is used to visualize the neuropil and Repo (magenta) to label all glia. White dashed boxes delineate the area magnified in the panels below. *10C12*-Gal4 is expressed specifically in epithelial and marginal glia, as well as in Lamina wide field neurons. *54H11*-Gal4 labels epithelial and marginal glia and medulla neuropil glia. *Hand-T2A*-Gal4 expression is only observed in some medulla neuropil glia (filled arrowheads), and absent in others (empty arrowheads). **(F-H)** FLEXAMP memory cassette is used to immortalize L3 expression of the Gal4 lines in **C-E** and to observe the morphology of adult glia. Only UAS-myrGFP expression is shown, and the neuropil shape is delineated in yellow dashed lines. Lamina and close-ups of medulla neuropil are indicated. Note that epithelial and marginal glia, that are indistinguishable at L3, diverge into specific morphologies and localizations in the adult **(F** and **G)**. Tracings of L3 medulla neuropil glia show that they give rise to all morphotypes (ensheathing and astrocyte-like) and localizations (proximal, lateral and distal) of adult medulla neuropil glia **(G** and **H)**. **xgo** = outer chiasm glia, **mng** = medulla neuropil glia, **lng** = lamina neuropil glia, **eg** = epithelial glia, **mg** = marginal glia. Gal4-UAS (*>>*). Scale bar = 20 µm.

We observed three distinct types of developmental trajectories of glial cell types based on our integrated dataset:

1. Cell types that did not exhibit a trajectory (*i.e.* minimally changed) from larva to adult, such as cortex and perineurial glia. These are cells that remain transcriptionally stable throughout development (**Fig. 4A**, black circles).
2. Cell types that gradually changed their transcriptome in a linear trajectory throughout developmental stages, like inner and outer chiasm glia, likely reflecting a maturation process (**Fig. 4A**, blue arrows). We also observed certain discrepancies between larval and adult cell types, which can be explained by previous literature. For example, in adults there are two satellite glia in the lamina named proximal and distal^16^. However, only one satellite glia is found in larvae, the larval satellite glia that gradually matures to become proximal satellite glia in adults. In contrast, the adult distal satellite glia has a different origin, corresponding to larval retina wrapping glia^23^, which is a larval nerve-wrapping glial type that is repurposed to work as cortex glia in adults. (**Fig. S2A, B**).
3. Cell types that formed an initial linear trajectory and then split into at least two different cell fates (**Fig. 4A**, red arrows). Notably, all glial types with a divergent trajectory were neuropil-associated glia that split from a common larval identity to adult **astrocyte-like (ALG)** and **ensheathing (EG)** cell types.

### Specification and diversification of lamina and medulla neuropil glia

In the adult **lamina neuropil**, there are two types of neuropil glia, known as epithelial glia (eg, a type of astrocyte-like glia) and marginal glia (mg, a type of ensheathing glia). eg/mg originate in the larva from specialized precursors localized in a region of the OPC called glia precursor center (GPC)^45^. At this stage eg and mg cannot be separated transcriptionally into distinct cell types, but later split into separated trajectories (**Fig. 4A**, **S2A, B**). We further investigated this split by UMAP trajectory analysis (**Fig. S3A**) and found that eg and mg transcriptomes diverge soon after they are born, i.e. between L3 to P15 (**Fig. S3B, C**). This apparent split into distinct cell types early during metamorphosis would explain the expanded cell diversity in the adult as compared to the larval scRNA-seq annotations. To confirm this trajectory, we performed memory-tracing experiments using the FLEXAMP memory cassette^55^, inducing recombination at late L3 using *10C12*-Gal4, a driver expressed in epithelial and marginal glia (eg/mg) as well as in Lamina wide-field (Lawf) neurons^45^ (**Fig. 4C**). In the adult brain, we recovered both epithelial (eg) and marginal glia (mg), confirming that lamina neuropil glia diverge into distinct cell types during pupal development (**Fig. 4F**). This result was independently confirmed using FLEXAMP with the *54H11*-Gal4 driver, which is expressed in eg/mg and medulla neuropil glia (see below).

The adult **medulla neuropil** presents two different neuropil glia subtypes, medulla ensheathing glia (EG, which have long projections closely associated with photoreceptors that show limited branching) and medulla astrocyte-like glia (ALG, that send dense, highly branched fine projections)^16^. Both EG and ALG can present diverse morphologies and positions around the neuropil, defined previously as ‘morphotypes’^16,17^. Both types derive from larval medulla neuropil glia (mng), which originate from the last division of neuroblasts in the outer proliferation center (OPC)^56,57^ and present a single trajectory that remains transcriptionally homogeneous from larva to P30 and then split into two different trajectories leading to the EG and ALG fates (**Fig. 4A, B**, and **Fig. S2A, B**). To validate this predicted trajectory, we induced FLEXAMP recombination in L3 using the *54H11*-Gal4 driver, which is expressed in medulla neuropil glia (mng) and eg/mg glia in larva (**Fig 4D**). Besides eg/mg lamina glia, we were able to recover adult medulla neuropil glial cells corresponding to both expected subtypes, EG and ALG glia. Furthermore, all adult morphotypes were observed, including distal and lateral medulla EG, as well as distal, lateral, and proximal medulla ALG^16,17^ (**Fig. 4G**). To further validate this result, we used another driver, *Hand-T2A*-Gal4. Although this driver is highly specific for L3 mng, not all larval mng were labeled by GFP (**Fig. 4E**). This sparsity was corroborated by our single cell analysis, where only 26% of L3 mng expressed *Hand* (**Fig. 4B**). We used a *memory-lacZ* cassette^58^ to trace L3 mng expressing *Hand-T2A*-Gal4 in the adult and found that *Hand* expressing cells were all Repo+ and localized around the medulla neuropil, consistent with the position of adult medulla neuropil glia (**Fig S2C**). We used FLEXAMP to assess their morphology and were able to identify both medulla EG and ALG cells of all morphotypes^16,17^. These results confirm that L3 mng initially share a common transcriptional identity and diverge into distinct EG and ALG subtypes during pupal development. These findings in lamina and medulla glia support the inferred bifurcation in the single-cell trajectory analysis.

### Molecular analysis of medulla neuropil ALG and EG diversification

To further analyze the basis of this transcriptomic divergence, we examined the trajectory of mng which splits into the medulla ALG and EG subtypes during pupal development. To identify the exact developmental stage of the split, we manually subset the cells at the fork of the trajectory and found that these cells were mostly coming from the P30 dataset (**Fig. 5C**, inset), suggesting that transcriptomic divergence of mng occurs much after their birth, unlike the lamina glia that split shortly after L3 (compare **Fig. 5A-D** with **S3A-C**). We then embedded a pseudotime trajectory using Monocle3^59^ on the existing UMAP reduction, which allowed us to closely recapitulate the developmental stages of the dataset as a function of pseudotime (**Fig. 5B, D** and see **Methods**). Since transcription factors (TFs) are the main drivers of cell fate, we focused on their expression to explore ALG and EG fate divergence. We found 63 TFs that were differentially and significantly modulated along the pseudotime trajectory (**Fig. 5E**), and we then used K-means to cluster TFs enriched at early, mid and late pseudotime points (see **Methods**). We found that TFs known to be required for mng specification, such as *gcm* and *gcm2*, as well as *Hand*, were expressed early in development, as expected^42,57^. Interestingly, we also observed early expression of canonical Notch effector genes such as *Hey* and five *E(spl)* complex TFs^60,61^, strongly suggesting that mng have active Notch signaling early after specification. This *Notch^ON^*status and initial *Hey* expression is also shared by lamina neuropil glia^45^. We validated the expression of *Hey* in eg/mg and mng by immunostaining, which showed that the presence of Hey protein in nascent lamina and medulla neuropil glia is transient and disappears soon after mng migration (**Fig. 5F, F’**). K-means clustering of TFs along pseudotime of the entire trajectory revealed that four genes, *crc*, *Eip78C*, *gem* and *CG3328* were enriched at the bifurcation point of ALG and EG subtypes, providing a list of candidate TFs that might be involved in mng subtype specification. Furthermore, we expanded this candidate TF list by analyzing the ALG and EG trajectories separately (**Fig. S3D, E**) which will serve as the basis for future perturbation-based experiments. Overall, our analysis provides a transcriptomic basis for mng subtype specification during optic lobe development.

**Figure 5.**
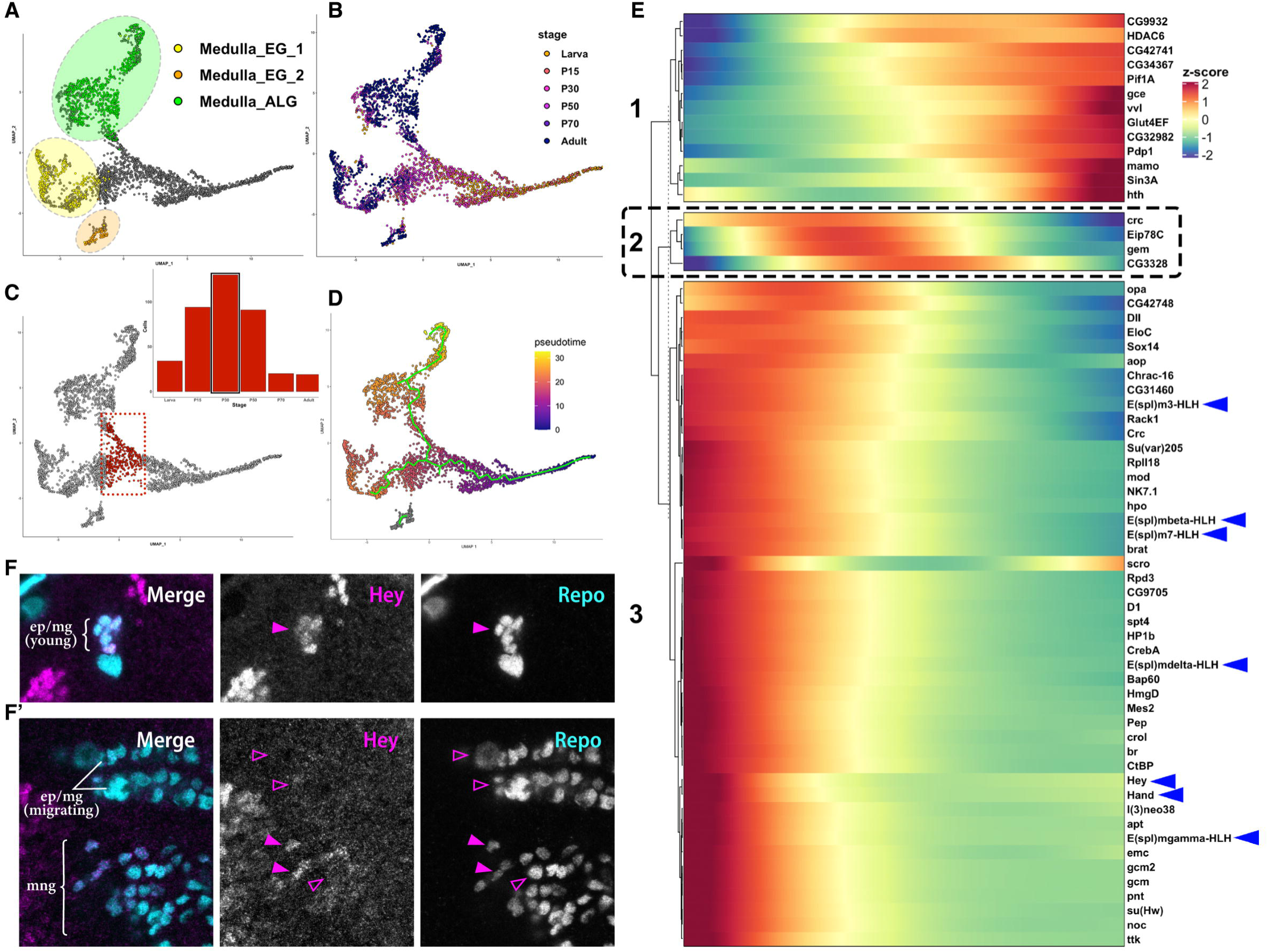
Transcriptomic Divergence of Medulla Neuropil Glia Along Development. **(A)** UMAP of medulla neuropil glia (mng) cells showing the divergence of astrocyte-like (ALG) and ensheathing (EG) glial cell types. The terminal fates identified in the adult dataset are circled. **(B)** UMAP showing cells colored by their developmental stages of origin. **(C)** UMAP-based subsetting of cells at the fork (colored red) between ALG and EG adult fates. Inset displays the proportion of red cells that belong to each developmental stage. Note that the most represented stage at the trajectory fork is P30. **(D)** UMAP showing pseudotime alignment of mng glial cells. This pseudotime embedding is used for analyzing gene expression dynamics in **(E)**. **(E)** Heatmap of dynamically regulated transcription factors along pseudotime highlighting putative TFs associated with mng divergence (dashed box) and canonical Notch effector genes (blue arrowheads). **(F, F’)** L3 brain showing the lamina **(F)** and medulla **(F’)** neuropil glia stained for Hey (magenta) and Repo (cyan) antibodies. Both lamina and medulla neuropil glia express Hey after specification (full arrowheads) and turn it off shortly after (empty arrowheads). **eg** = epithelial glia, **mg** = marginal glia and **mng** = medulla neuropil glia.

### Specification of lobula neuropil glia

The glia associated with the **lobula complex neuropil** (which during pupal development forms the **lobula** and **lobula plate** neuropils) seems to develop in a more complex manner than what we found for medulla and lamina neuropil glia.

We annotated and validated one cluster as lobula complex neuropil glia in the larva (**Fig. 2A, B** and **I**). However, unique markers to validate lobula neuropil glia in adults were not found (**Fig. 3A**). Since we confidently annotated all remaining glial types, the 6 unannotated glial clusters in the adult were good candidates to be lobula neuropil glia. Moreover, these unknown adult clusters were closely associated with the L3 lobula complex neuropil glia cells in our age-integrated dataset (**Fig. 4A**). To gain further insight into the presumptive lobula neuropil glia, we integrated these clusters across developmental stages (**Fig. S2E, F**). We named the presumptive ALG clusters “Astrocytes_1/ 2/ 3/ 4” and presumptive EG clusters “Ensheating_1/ 2/” (**Fig. S2F**). We observed two parallel trajectories: one for lobula neuropil ALG, which originated in the validated L3 lobula complex neuropil glia cluster, and another for lobula neuropil EG, that started at P15 (**Fig. S2F**).

Interestingly, the L3 lobula complex cluster already expressed mature astrocyte-like glia markers and was positioned between adult medulla ALG and the unknown adult astrocyte-like clusters in the integrated data UMAP (see “Lobula C. Neuropil” and “Astrocytes_1/ 2/ 3/ 4”, **Fig. S2A, B**). This suggests that the lobula complex neuropil glia found in the larva are already mature, astrocyte-like cells. Intriguingly, the lobula complex is physically closely associated with the central brain during larval development^19^. In the central brain, adult (or secondary) neuropil glia is produced during larval stages from type II neuroblasts to replace larval (or primary) neuropil glia born during embryogenesis, raising the possibility that the L3 lobula complex neuropil glia cluster was composed of mature astrocyte glia migrated from the central brain. Accordingly, we have observed that a population of glial cells surrounding the lobula complex neuropil that expressed *54H11*-Gal4 appeared to migrate from the central brain (**Fig. S2D)**. It is possible that these cells account for the lobula complex neuropil cluster of larval origin that is at the base of the lobula neuropil ALG trajectory (**Fig. S2E**, yellow cells).

Meanwhile, the presumptive lobula neuropil EG trajectory, which gives rise to “Ensheathing_1/ 2/” adult clusters, started at P15 and did not include any L3 cells (**Fig. S2E, F**). We looked for genes that were expressed early in both medulla and lamina neuropil glia developmental trajectories, and found 61 common genes (**Table 3**), six of which were unique for these clusters and mostly absent in other glial types: *CR30009, gcm, E(spl)mgamma-HLH, CG11670, E(spl)malpha-BFM* and *Traf4* (**Fig. S4A-F**). Interestingly, these genes were also expressed at the beginning of the lobula neuropil EG trajectory but were almost absent from the lobula neuropil ALG trajectory (**Fig. S4G-I**). This observation supports the idea that ALG lobula neuropil glia is more mature than the rest of neuropil glia at L3 and suggests that EG lobula neuropil glia are born at later stages (P15).

Finally, there were three clusters of surface lamina glia for which we could not identify contributions from larva glia: pseudocartridge (a subtype of subperineurial glia) as well as fenestrated and chalice glia, which are both subtypes of perineurial glia (**Fig. 4A** and **S2A, B**, dashed circles)^16^. Pseudocartridge, fenestrated glia and likely chalice glia originate from the eye disc^62^ that was not sequenced in the larva dataset^25^, thus explaining their absence (summarized on **Table S3**).

Together, our data show that the increased number of adult glial clusters compared to the larval dataset is, in part, the result of increased glial cell diversity originated from larval neuropil glia that diverges into different subtypes during pupal development.

### scRNA-seq reveals subcellular distribution of RNAs

Next, we sought to investigate three surprising observations in the larval dataset:

- First, we found a cluster corresponding to retina wrapping glia in our larval dataset (**Fig. 2B**, and **K**), which was unexpected, since their cell bodies are located in the eye disc and optic stalk^63,64^ that were dissected out from the larval optic lobe before sequencing^25^ (**Fig. 6A**). In contrast, the other eye disc-derived glial types (chalice, pseudocartridge, and fenestrated, see above) were absent. Interestingly, the retina wrapping glia are the only eye disc glia that extends their processes into the optic lobe. Therefore, we hypothesized that cell processes that were broken off the cell body could form fragments that were sequenced and contain enough mRNA to form clusters of a defined cell identity.

**Figure 6.**
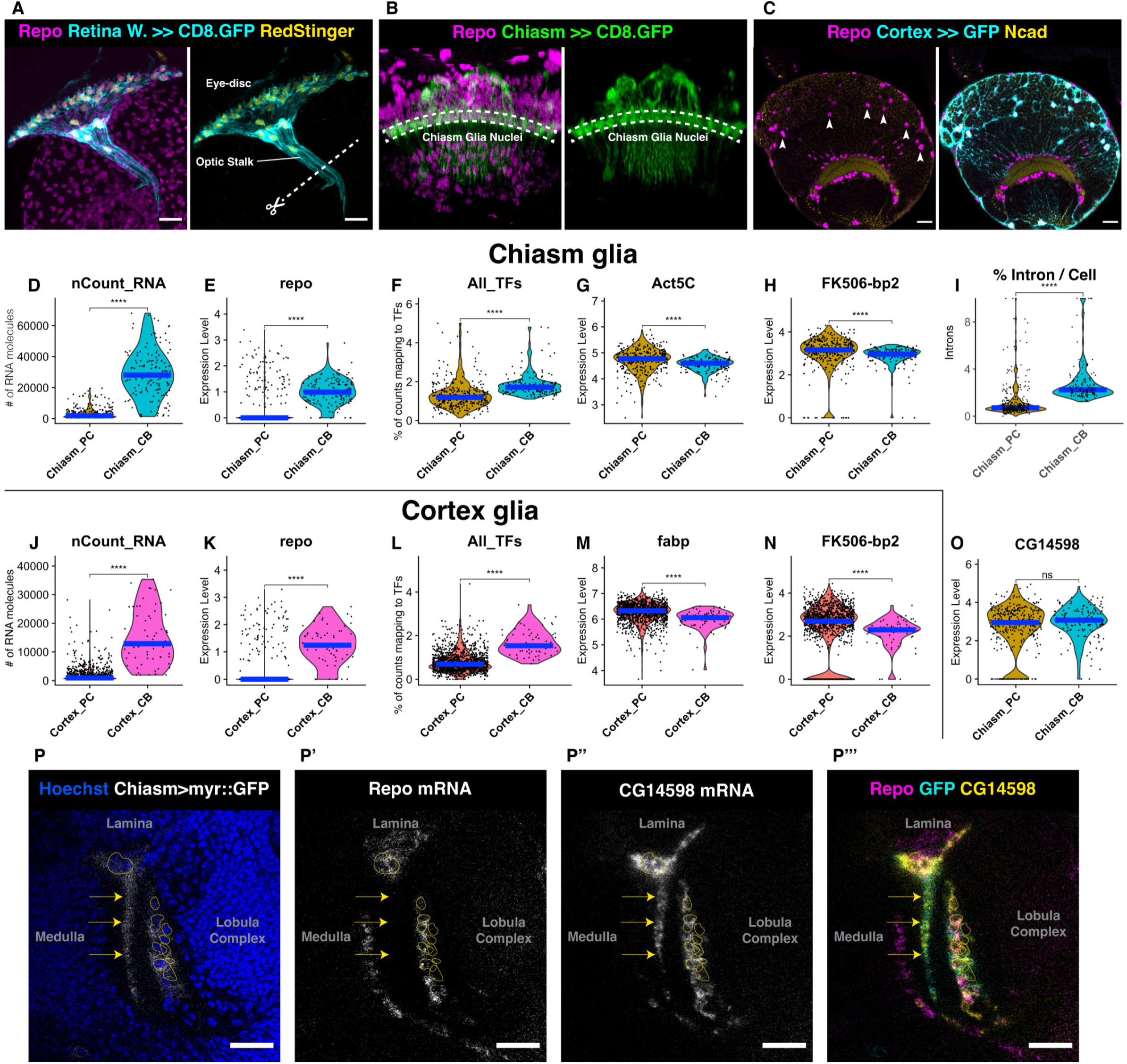
scRNA-seq Reveals Subcellular Distribution of RNAs. **(A)** Retina Wrapping glia Gal4 driving expression of membrane-bound GFP (cyan), RedStinger (nuclear, yellow) and Repo (magenta) to identify glial cells. The dashed line shows where the optic lobe and eye-disc were separated during dissection prior to the sequencing of the optic lobe. **(B)** Chiasm glia Gal4 driving expression of membrane-bound GFP (green) and Repo antibody (magenta). The white dashed line shows the position of chiasm glia cell bodies. **(C)** Cortex glia-Gal4 (*Cyp4g15*-Gal4) driving expression of GFP (cyan). Repo antibody (magenta) and Ncad shows the position of the medulla neuropil (yellow). Some cortex glia somas are indicated by a white arrowhead. **(D-O)** Violin plots comparing Chiasm_CB and Chiasm_PC **(D-I** and **O)** or Cortex_CB and Cortex_PC **(J-N)**. Statistics by Wilcox-test, p-value < 0.00005 = ****, p-value > 0.05 = non-significant (ns). The blue bar represents the median. **(D)** Number of UMIs per cell. Chiasm_CB have significantly more UMIs than Chiasm_PC. **(E)** Expression level of repo. Chiasm_CB contains significantly more repo than Chiasm_PC. **(F)** Percentage of reads that align to all transcription factors. Chiasm_CB have significantly more reads than Chiasm_PC. **(G)** Expression level of *Act5C*. Chiasm_PC contains significantly more *Act5C* than Chiasm_CB. **(H)** Expression level of *FK506-bp2*. Chiasm_PC contains significantly higher *FK506-bp2* than Chiasm_CB. **(I)** Percentage of valid reads mapping to introns. Chiasm_CB has significantly more introns than Chiasm_PC. **(J)** Number of UMIs per cell. Cortex_CB has significantly more UMIs than Cortex_PC. **(K)** Expression level of *repo*. Cortex_CB contains significantly higher *repo* than Cortex_PC. **(L)** Percentage of reads that align to all transcription factors. Cortex_CB has significantly more reads than Cortex_PC. **(M)** Expression level of *fabp*. Cortex_PC contains significantly higher *fabp* than Cortex_CB. **(N)** Expression level of *FK506-bp2*. Cortex_PC contains significantly higher *FK506-bp2* than Cortex_CB. **(O)** Expression level of *CG14598*. Both clusters have similar levels of *CG14598*. **(P)** HCR RNA FISH of the L3 optic lobe expressing membrane-bound GFP in chiasm glia. Yellow outlines highlight chiasm glia nuclei. Arrows point to chiasm glia processes. Hoechst shows nuclei of all cells (blue). *DAT*-Gal4 drives membrane-bound GFP (grey) expression in chiasm glia. *GFP* mRNA was detected by FISH. **(P’)** *repo* mRNA is found around the nuclei, but not in the processes of chiasm glia. **(P’’)** *CG14598* mRNA is found around the nuclei and in the processes of chiasm glia. **(P’’’)** Merged image, *repo* (magenta), *GFP* (cyan) and *CG14598* (yellow). Scale bar = 20 µm.

- Second, we found an extra chiasm glia cluster that could also be explained by the sequencing of cellular processes since this glial type also presents long and extensive cellular processes (**Fig. 6B**). At larval stages, unsupervised clustering revealed two chiasm glia clusters: Cluster 12 that could be separated with supervision and annotated as inner and outer chiasm (**Fig. 2B** and **Fig. 7A**), the only two known types of chiasm glia in the optic lobe^65^. On the other hand, cluster 4 shared all the top chiasm glia markers with cluster 12 (**Fig. S5B**), suggesting that cluster 4 was not a different third type of chiasm glia. We thus hypothesized that cluster 12 was composed of <u>c</u>ell <u>b</u>odies with nuclei (Chiasm_CB), while cluster 4 was comprised of <u>p</u>ro<u>c</u>esses without nuclei (Chiasm_PC) (**Fig. S5A, B**).

**Figure 7.**
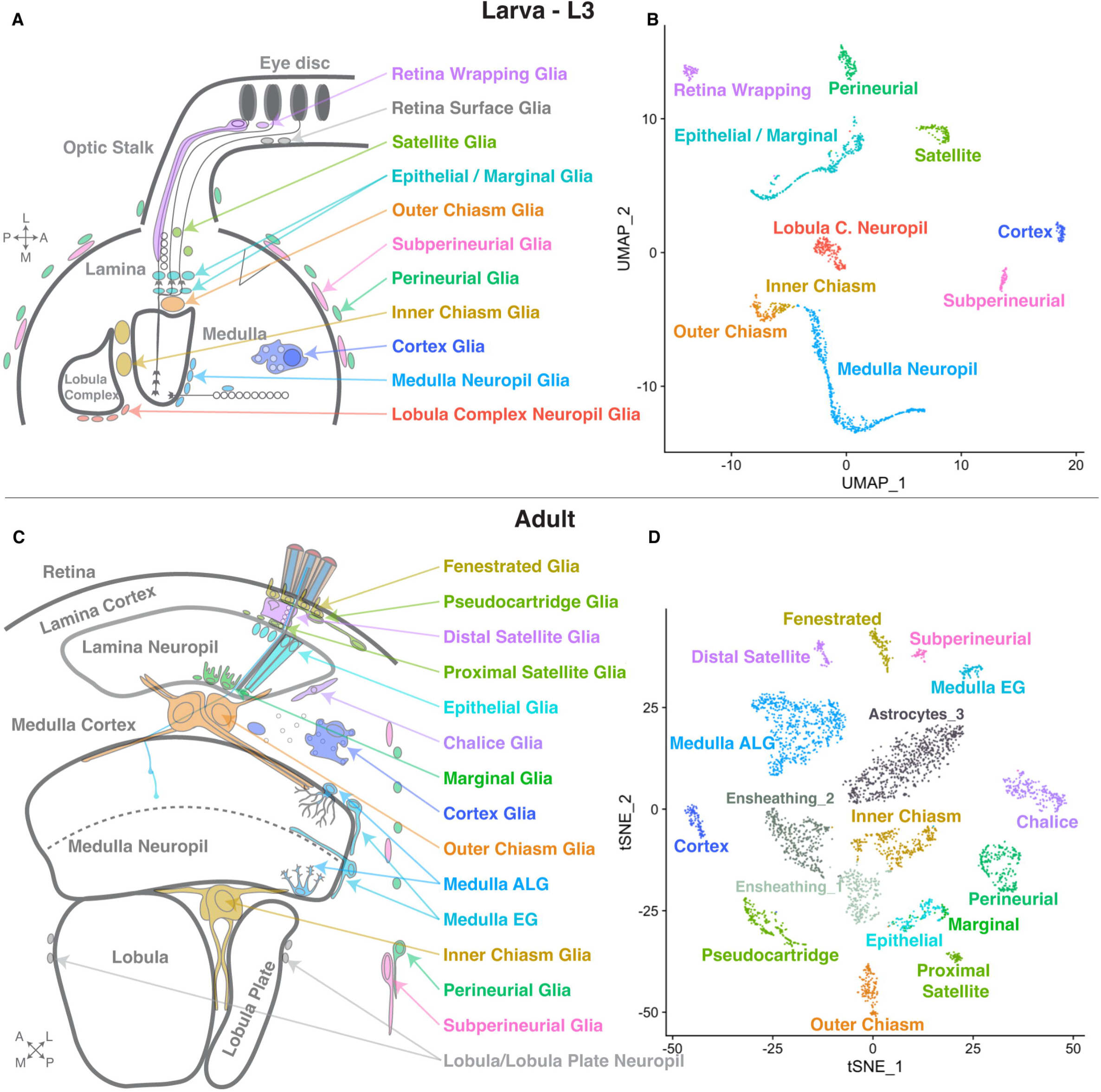
Annotation of Optic Lobe Glia in the Larval and Adult Single-Cell Datasets. **(B)** A cross-section schematic representation of the larva (L3) optic lobe with the glial cell types depicted. **(C)** UMAP visualization of the final larval glial cells annotations after removal of cellular processes in the dataset (**Table 1**). Colors of the clusters match the colors in the drawing in **(A)**. **(D)** A cross-section schematic representation of the adult stage optic lobe with the glial cell types depicted. **(E)** TSNE visualization of the final adult stage glial cells annotations after removal of cellular processes in the dataset (**Table 2**). The colors of the clusters match the colors in the drawing in **(C)**.

There were two striking differences between the putative chiasm cell body and the putative chiasm processes clusters. First, the Chiasm_CB cluster contained cells with significantly more molecules of RNA (UMIs) than Chiasm_PC (**Fig. 6D**); second, only Chiasm_CB contained mRNA for the transcription factor (TF) *repo* (**Fig. 6E**) and the combined content of all TFs was significantly higher in Chiasm_CB compared to Chiasm_PC (**Fig. 6F**). In contrast, Chiasm_PC presented higher levels of *Act5C* (**Fig. 6G**) and *FK506-bp2* (**Fig. 6H**) as well as other genes previously described as having a specific sub-cellular localization in cell extensions (**Table S2**). For example, the closest human ortholog of *Act5C* is β*-actin*, the first gene to be shown to have its mRNA subcellularly localized and transported to the neurites of neurons and lamellipodia of migrating fibroblasts^66–69^. *FK506-bp2* is a member of a highly conserved family of immunophilins that are also enriched in neurites in mammalian neurons^70^. To strengthen our hypothesis, reads mapping to introns or exons were compared in both chiasm clusters. Since unspliced mRNAs and introns are essentially only present in the nucleus, Chiasm_CB should contain relatively more reads mapping to introns than Chiasm_PC. Indeed, Chiasm_CB contained, on average, 3% of reads mapping to introns versus 1.3% in Chiasm_PC (**Fig. 6I**).

To confirm that non-nucleated cell fragments were generated during our cell dissociation protocol, we dissociated the optic lobes of flies expressing membrane-bound GFP in chiasm glia following the same protocol used to generate the larva scRNA-seq dataset^25^. We observed similar-sized cell fragments of GFP-expressing chiasm glia that contained (**Fig. S5C**, top panel) or did not contain (**Fig. S5C**, lower panel) nuclei stained with Hoechst.

- Third, we sought to investigate if cell fragmentation could explain the high numbers of cortex glia observed in our scRNA-seq dataset, which was much larger than expected. By nuclear counting using an anti-Repo antibody, we estimated that cortex glia represents less than 7% of all glia in the larval optic lobe (about 100 cortex glia out of about 1500 glia per optic lobe). However, they comprised the largest cluster in the larval dataset – about 43% of all optic lobe glial cells (1332 out of 3117 cells, after excluding central brain glia) (compare **Fig. 6C** and **S5A, B**). Like for chiasm glia, cortex glia clusters could be separated into cell bodies (Cortex_CB) and processes (Cortex_PC). Both clusters were transcriptionally similar (**Fig. S5B**), but Cortex_CB contained many more UMI, a higher amount of *repo* mRNA and more reads encoding TFs than Cortex_PC (**Fig. 6J-L**). Cortex glia are very large cells that have processes surrounding the somas of all cells, including neurons, neuroblasts, and the neuroepithelium^16^ and might therefore be susceptible to break into multiple fragments by the cell dissociation protocols used to generate scRNA-seq datasets (**Fig. 6C**). This likely explains the high number of Cortex_PC found in the dataset (41% of all glial “cells” in the L3 dataset – 1270 out of 3117 cells) compared to Cortex_CB (2% - 62 out of 3117 cells). Cortex_PC expressed higher levels of *fabp* and *FK506-bp2* than the Cortex_CB cluster (**Fig. 6M, N**), among other genes (**Table S2**). Fabp is a fatty acid binding protein whose mouse ortholog *Fabp7* mRNA is localized to the tripartite synapses in astrocytes and coimmunoprecipitates with the cytoplasmic polyadenylation element-binding 1 (CPEB1), which is responsible for regulating subcellular mRNA trafficking and translation inside cells^5,71^. In conclusion, the sequencing of cellular processes without nuclei appears to explain the large discrepancy in the number of cortex glia cells in the dataset.

Finally, we sought to determine whether there was indeed a differential subcellular distribution of mRNAs in chiasm glia *in vivo*. We performed HCR RNA-FISH for *repo* (Chiasm_CB only) and used as a control *CG14598*, a gene that is highly and specifically expressed in chiasm glia but not significantly enriched in cell bodies or processes (**Fig. 2A** and **6O**). In a fly line expressing membrane-bound GFP in chiasm glia to identify the location of cell processes, *repo* mRNA was only found around the nucleus in the cell body, while *CG14598* mRNA was found throughout the cell (**Fig. 6P**), which was consistent with the Chiasm_CB cluster corresponding to the cell bodies and Chiasm_PC to the cell processes.

Taken together, these results show that when large cells such as certain glia are dissociated with commonly used scRNA-seq protocols, pieces of cells can form “pseudo cells” that can be mistakenly identified as distinct cells using classical scRNA-seq clustering algorithms. This phenomenon explains the presence of retina wrapping glia, the third cluster of chiasm glia, and the overrepresentation of cortex glia in our larval dataset.

### Computational approaches to detect cellular processes in scRNA-seq datasets

We aimed to comprehensively identify and eventually remove cellular processes from our datasets to build an accurate atlas of the developing optic lobe glial cells (**Fig. S6A**). We have previously established that droplets containing cellular process fragments had significantly fewer mRNAs while exhibiting an unambiguous transcriptomic similarity to the clusters annotated as their corresponding cell bodies, and that they were also relatively deprived of TF-encoding transcripts. Therefore, a more robust dataset could be generated by identifying and excluding cellular processes, which are the source of overrepresentation of some glia and of extra clusters in the datasets.

We used two approaches to identify potential cellular process-containing clusters. These clusters contain lower total UMIs and lower TF-encoding reads when compared to either the whole dataset (approach 1) or when compared to the most similar cluster with high UMIs based on Pearson correlation (approach 2) (**Fig. S6A**).

Using both approaches for the larval dataset, we confirmed the previously identified clusters of chiasm and cortex glia that were investigated above as being cellular processes (**Fig. S6B**). Moreover, approach 1 also detected retina wrapping as processes and a newly identified cluster of processes of subperineurial cells. However, because these clusters were the only ones of their identity, they were not removed from the dataset. Moreover, both approaches identified two transcriptionally similar clusters of perineurial glia, one of them likely being the cellular processes of the other. Similarly to retina wrapping, chiasm and cortex, perineurial glia extend large cellular processes that extend around the whole optic lobe^72^. In summary, for the larval dataset, we excluded clusters comprised of cellular processes for chiasm, cortex, and perineurial glia.

In contrast to the larval dataset where cell types were subdivided into multiple clusters, the adult dataset had most of its cell types identified as unique clusters; therefore, to better understand the possible presence of cellular process clusters in this dataset we over-clustered the dataset into smaller subclusters and used both approaches described before (**Fig. S6C-E**). Both approaches showed that clusters Astrocytes_1, Astrocytes_2, and Astrocytes_4 were likely entirely comprised of cellular processes. A subcluster of Ensheathing_1 also showed up as processes in both approaches. Adult perineurial glia could also be divided into cell processes and cell bodies, and the same was observed for cortex, distal satellite and medulla ensheathing glia. In summary, we excluded 3 clusters of astrocyte-like glia, 1 subcluster of ensheathing glia, 1 cluster of cortex glia, 2 subclusters of perineurial glia and 1 subcluster of distal satellite glia (**Fig. S6**).

Taken together, these different approaches identified cellular processes that were the source of extra clusters in the dataset. This allowed us to build a more accurate atlas of the developing optic lobe glial cells in the larval (**Fig. 7A, B**) and adult stages (**Fig. 7C, D**) and assess the transcriptome of only the cell bodies (**Table S1**).

## Discussion

Neuronal development highly depends on glial function^1–4^. To study glial development and diversity and its contributions to brain formation and function, it is essential to have access to its annotated transcriptome throughout development. We comprehensively annotated all the optic lobe glial clusters from larva to adult stages using previously generated scRNA-seq datasets and collected marker gene information for each cluster (**Table S1**). Combined with our previous developmental scRNA-seq neuronal datasets^24^, this work provides a powerful tool to study neuron-glia interactions and glial development and function.

*Repo* and *MRE16* (also known as *CR34335*) long-noncoding RNA were previously proposed to be useful in identifying glial clusters^52^. However, while Repo protein is a very specific marker for glial cells^26^ and it is the standard to identify glial cells by immunostaining, we showed here that its transcripts are lowly expressed and are often not detected in glial clusters in scRNA-seq datasets. In contrast, *MRE16* is not specific to glial cells and is not a reliable marker, as observed for a VNC dataset that used both *repo* and *MRE16* and could not confidently annotate all glial clusters based on these markers, alone^38^. To overcome these problems, we found three additional markers *CG32032*, *AnxB9* and *GstE12*, and have developed antibodies for AnxB9 and CG32032 proteins. We were able to confirm the specificity of the AnxB9 antibody by RNAi knockdown, but could not do so for CG32032 antibody, probably due to inefficacy of the available RNAi construct (**Fig. S7**). However, the knockdown of *AnxB9* in outer chiasma (xgo) and wrapping glia (wg), did not appear to affect these cell types, and more work will be required to understand the functional role of these markers in glia development. Importantly, the expression of these markers is not necessarily restricted to glia, as exemplified by the expression of *AnxB9* in VNC neurons. Thus, we propose to use the combination of two or more of these markers, together with or independently of *repo*, to faithfully identify all glial clusters in any given *Drosophila* scRNA-seq dataset.

After annotating the glial clusters in larva and adult, we were intrigued by the increased glial diversity that we observed in adult stages as compared to the larva, and we decided to leverage our developmental transcriptomic atlases to investigate this phenomenon. With this approach, we proposed and validated different “developmental trajectories” through which some glial types remained transcriptionally stable, matured, adapted or diversified during pupal development. Some cells like cortex and perineurial glia did not form a trajectory, and cells from different developmental stages were transcriptionally identical. This was expected, since cortex and perineurial glia are generated before the beginning of optic lobe differentiation and are therefore already specified and mature when the larval cells were sequenced^62,72–74^. Moreover, their functional roles, trophic support for cortex glia and barrier formation for perineurial glia, change little through development, explaining a stable transcriptomic profile across stages.

Most glial types, however, displayed transcriptional changes from larval through adult stages. For instance, inner and outer chiasm glia formed a trajectory through development, gradually changing to acquire their final fate in adulthood. These transcriptional changes occurred gradually in a straight trajectory, likely reflecting a process of maturation, process extension, etc. In contrast, retina wrapping glia changed their functional type from nerve-wrapping in the larval retina to cortex glia (distal satellite glia) in adults. Retina wrapping glia coordinate neuronal differentiation and axon guidance during larval stages^3,75^, and serve to protect neurons crossing from the eye disc to the optic lobe. However, because the retina directly abuts the lamina in adults, there is no longer need for wrapping glia around photoreceptor axons between these structures. Our data suggests that these glial cells do not die and are instead repurposed to serve as cortex glia.

Most interestingly, we found that lamina and medulla neuropil glia initially followed a single trajectory that later split into two branches during pupal stages to achieve different fates in adults. An interesting example of this trajectory is medulla neuropil glia, which is produced during the last division of medulla neuroblasts, after they have given rise to different medulla neurons^56,57,76^. Medulla neuropil glia migrate towards the neuropil over the surrounding neuronal somas and place their cell bodies closest to the medulla neuropil^20^. At this larval stage, mng did not seem to have acquired a terminal fate, and our data showed that at this point cells were mostly homogeneous at the transcriptional level. However, they start exhibiting differential gene expression after P30 when two subtypes could be observed: astrocyte-like (ALG) and ensheathing glia (EG). Interestingly, branch extension of ALG mng, a cell autonomous process dependent on the transmembrane leucin-rich repeat protein Lapsyn and the FGF receptor Heartless (Htl) starts around P60, after our proposed mng trajectories diverge^20^. Notably, *lapsyn* and *htl* loss of function cause unbranched mng that resemble the morphology of adult EG mng ^16,17,20^. It would be interesting to analyze whether the differentially expressed TFs in the ALG mng trajectory found in this study regulate *lapsyn* and *htl* expression to achieve the branched morphology characteristic of astrocyte-like glia.

Three hypotheses might explain how neuropil ALG *vs.* EG final fate is determined: 1) the fate is determined at birth with undetectable transcriptional differences; 2) the fate is influenced by external cues depending on their final location in the neuropil, or 3) cell fate is stochastically determined later. We observed that the transcription factor *Hand* was expressed in only a fraction of mng in L3, which could account for early ALG vs EG fate differences. However, our lineage tracing experiments demonstrated that *Hand*-expressing mng give rise to both astrocyte and ensheathing morphologies, arguing in favor of a single mng glial type before P30. ALG and EG lamina neuropil glia display a clear spatial segregation in the adult: epithelial glial cells (ALG) are located in the distal lamina, whereas marginal glial cells (EG) are in the proximal end of the lamina neuropil^16,17,45^. Interestingly, two rows of lamina neuropil glia (one at the proximal and the other at the distal side of the developing lamina plexus) are readily observed at L3^42^. However, at L3 lamina neuropil glia (epithelial and marginal) appeared transcriptionally identical, only showing clearly separated clusters around P30. Previously, live-imaging of these glia in larvae showed that young eg/mg cells can interchange their position crossing through the lamina plexus, exemplifying their plastic identity at L3 stage^77^ (and our unpublished data). Based on this evidence, we hypothesize that lamina eg and mg fates are not determined at birth but are established later during early pupal development based on their final location on either side of the growing lamina neuropil, probably in response to local cues. By contrast, ALG and EG mng are located next to each other surrounding the whole perimeter of the adult medulla neuropil, making it difficult to propose that external signals determine their final fate. In this case, the more likely scenario is that a stochastic decision triggers ALG- or EG-specific genetic programs. It has been recently shown that all astrocyte-like glia in the optic lobe are remarkably homogeneous transcriptionally, regardless of their multiple morphologies (morphotypes) or locations^17^. Consistently, we have recovered every morphotype of mng from our larval FLEXAMP tracings. Therefore, our study confirms that both lamina and medulla neuropil glial types are born as a single identity that bifurcates during pupal development into astrocyte-like and ensheathing glia through different mechanisms.

To our knowledge, the origin of lobula complex neuropil glia remains to be elucidated. However, the closest glia associated with them in our larval hierarchical clustering are chiasm glia which derive from central brain neuroblasts^19,78^. We also observed a significant number of glial cells expressing *54H11*-Gal4, a driver shared with lamina and medulla neuropil glia that seem to migrate from the central brain to the lobula complex. Moreover, a recent scRNA-seq dataset that sequenced cells derived from medulla precursor cells found glial cells corresponding to the clusters we identified as medulla neuropil ALG and EG glia, but not lobula neuropil glia (Astrocytes_3, Ensheathing_1 and Ensheathing_2)^79^. All this evidence leads us to hypothesize that lobula neuropil glia derive from central brain progenitors. The central brain develops earlier than the optic lobe and thus has mature astrocyte-like glia in L3^19,80^, which could explain the presence of markers associated with mature astrocyte-like glia in the lobula complex neuropil glia cluster we have validated in L3 that stands at the root of adult lobula neuropil ALG clusters. In contrast, lobula neuropil EG glia can only be identified from P15 onwards, express markers related to early development in other neuropil glia and seem to display a parallel trajectory to lobula neuropil ALG rather than derive from them. Therefore, we hypothesize that lobula neuropil ALG and EG have a central brain origin, albeit they are born and migrate towards the lobula complex at different developmental times. However, specific experiments to test this hypothesis are still necessary.

Finally, we showed here that during cell dissociation large glial cells are fragmented into pieces that contain the nucleus (cell body) or not (processes). The cellular process fragments still hold enough RNA to be sequenced and annotated as unique cells and to pass through the quality control steps. Interestingly, the relatively fewer molecules of RNAs in processes is still sufficient to allow them to cluster with or next to their cell bodies. We devised approaches to identify cell process clusters by comparing the UMIs and transcription factor mRNAs within a cluster against the whole dataset or against their cell body counterpart. This allowed us to eliminate cell processes’ clusters and produced a more accurate glial atlas of the larval and adult stages optic lobe. However, identifying cell processes in a scRNA-seq dataset can become a powerful tool to study subcellular localization of mRNAs, and can become a technical advantage when studying cells that are hard to isolate and to physically separate the cellular compartments.

In summary, this work presents a comprehensive developmental transcriptomic atlas of optic lobe glia. By integrating larval-to-adult datasets, we uncover multiple developmental strategies–stability, maturation, repurposing, and fate bifurcation–through which glial cells achieve adult identities. Importantly, several neuropil glia arise as a single, plastic population that later diversifies into astrocyte-like and ensheathing subtypes. Finally, by identifying single cell transcriptomic clusters formed by cell processes, we refine atlas accuracy and highlight new opportunities to study subcellular RNA localization. Together, this work provides a foundational framework for investigating glial development and neuron–glia interactions in the *Drosophila* optic lobe.

## Methods

### Drosophila strains

Flies were kept at 25 °C 12-hour light/dark cycles on a standard cornmeal medium. 10xUAS-myr::GFP (BDSC #32197), UAS-RedStinger (#8546), Tret1-1-Gal4 (#67471), Vmat-Gal4 (#66806), rost::LacZ (#12042), CG3036::GFP (#50804), b-Gal4 (#76724), DAT-Gal4 (#84622), CG31663-Gal4 (#76708), ome-Gal4 (#76166), Canton-S (#64349), Hand-T2A-Gal4 (#66795), CG4288-Gal4::GFP (#29208), alphaTub85E::GFP (#60267), Npc2b-Gal4::GFP (#24694), Adgf-D-Gal4 (#77799), R25A01-Gal4 (#49102), UAS- CD8:GFP (from Bloomington), Cyp4g15-Gal4 (#39103), GstT4-Gal4 (#77777), nkd::LacZ (#25111), R94A08-Gal4 (#40673), R10C12-Gal4 (#47841), R54H11-Gal4 (#39094), UAS-AnxB9-RNAi (#38523), UAS-AnxB9-RNAi.GD (VDRC #27493), wg/xg^O^-Gal4 (#9488), UAS-CG32032-RNAi.GD (VDRC #5167), Act5C>stop>nuc.lacZ; UAS-Flp (memory-lacZ) (Struhl and Basler 1993), 20XUAS-Flp; Gal80ts; Act>y+>LexA, LexAop-myr:GFP (FLEXAMP) ^55^, UAS-sfGFPnlsPEST-2A-B2ase (UAS-nGFP, for short across the manuscript) – this is an unpublished line developed by Dr. Ben Jiwon Choi (Desplan lab), consisting of a destabilized nuclear-localized GFP that minimizes fluorophore perdurance. The complete genotype for each experiment can be found in the **Table Fly Genotypes**.

### Immunohistochemistry

Fly brains from adult stage or larva were dissected in room temperature 1x PBS. Then fixed in 4% formaldehyde (v/w) in 1x PBS for 20 minutes at room temperature. Following the fixation, the formaldehyde solution was removed, and the brains were washed 3x in PBSTX (0.5% Triton X-100 in 1x PBS). Then, the brains were blocked in block solution (PBSTX with 5% Horse Serum) for at least 30 minutes at room temperature. Primary antibody was diluted as specified in block solution and the samples were incubated for two days at 4 °C with primary antibody. Then the primary HCRantibody solution was removed, and the brains were rinsed 3x with PBSTX and 1x washed with PBSTX for at least 2 hours. Then secondary antibodies were diluted in the block solution and added to the samples, which were incubated for two days at 4 °C. Then, the samples were washed 2x with PBSTX and 1x with 1x PBS. Slides were mounted using a 0.22 mm spacer and samples immersed in Slowfade. Images were acquired in a Leica SP8 confocal and were processed in Fiji and Illustrator. The antibodies used are listed in (Table – antibodies).

### HCR-RNA FISH

To perform fluorescence in situ hybridization (FISH) for the detection of RNAs, we used custom probes designed for *Repo, CG14598, and Cpr51A*. All probes were designed by Molecular Instruments^29^. Hybridization buffer, amplification buffer, wash buffer, and fluorophore-labelled amplification hairpins (Alexa 546, 647, 594, and 488) were obtained from Molecular Instruments. We followed steps and reagents in the published protocol^81^. For combining HCR-RNA FISH and Immunohistochemistry, we performed the Immunohistochemistry first but using Molecular Instrument’s Antibody Buffer instead of PBSTX. Then fixed the tissue again in 4% PFA in PBST for 20 minutes at room temperature and then performed the HCR-RNA FISH protocol.

### Antibody generation

Polyclonal antibodies were generated by Genscript (https://www.genscript.com/). The protein sequences used for each immunization were as follows:

AnxB9 (guinea pig), MSSAEYYPFKCTPTVYPADPFDPVEDAAILRKAMKGFGTDEKAIIEILARRGIVQRLEIAEAFKTSYGKDLISDLKSELGGKFEDVILALMTPLPQFYAQELHDAISGLGTDEEAIIEILCTLSNYGIKTIAQFYEQSFGKSLESDLKGDTSGHFKRLCVSLVQGNRDENQGVDEAAAIADAQALHDAGEGQWGTDESTFNSILITRSYQQLRQIFLEYENLSGNDIEKAIKREFSGSVEKGFLAIVKCCKSKIDYFSERLHDSMAGMGTKDKTLIRIIVSRSEIDLGDIKEAFQNKYGKSLESWIKMSIKSLTYVAIFGLFWGSIAGTVVDQFGIYGGSPITTTERSNAELRCMNINPQN SVDLEQ;

CG32032 (rabbit), MMGLWYGSEIIVHSQDFPGTYEYDSCVIIHLTDATDQIRLSQANRGYGYGNQDYNRNQNNYGRTTTTQSSYPDSDEYPLRSIQSQQKYLRLIWSERDNNLEYTFNYTTSAPGQWSNIGDQRGSLVTLNTYTQFTGTVQVVKAVNDHLVLTFCGNDVKSSIYTVVLTRNRLGLSLDELRSIRNLLSRRGLYTETIRKVCNGCGRLGGSLFALLALLLVVRLAWGRGQ;

Optix (rabbit), AFSAAQVEIVCKTLEDSGDIERLARFLWSLPVALPNMHEILNCEAVLRARAVVAYHVGNFRELYAIIENHKFTKASYGKLQAMWLEAHYIEAEKLRGRSLGPVDKYRVRKKFPLPPTIWDGEQKTHCFKERTRSLLREWYLQDPYPNPTKKRELAKATGLNPTQVGNWFKNRRQ R;

Repo (rat), MEHDSFDDPIFGEFGGGPLNPLGAKPLMPTTTAMHPVMLGSVHELCSQQQQQQQQQRLPDCNTILPNGGGGGAGSGGAGGSPNYVTKLDFVNKMGCYSPSQKYEYISAPQKLV EHHHHHH;

Hey (rat), MDHNMHVNAPSLHHWGYAAGPGVVMPGATATTPQSHWVPPPQSHHSAHSNHSHGHSQGHSHGIGSLKRTLSESDCDDLYSEESSKEQISPSEPGSCQLMSRKKRRGVIEKKRRDRINSSLTELKRLVPSAYEKQGSAKLEKAEILQLTVEHLKSLQSKTLDSLSYDPQRVAMDYHIIGFRECAAEVARYLVTIEGMDIQDPLRLRLMSHLQYFVQQRELSAKSCASPGGWSPAAPSSSGYQPNCAAAPYQSYAAPANPGAYVSSYPTLSASPSQQAQQLGGRTSVSRTSGSAVTESLPSHDLHSDSSSQQQQQQQQQQQQQQQHQQQQHQQQQQRTQTT PQPTQQQHYTHDHSAVHSEQQVPTYIELTNSNRPAAIGSDSLSYSAAPQYPVSGLPGQDYNNSSVLQYATPNGAKPYRPWGAEMAYHHHHHH.

Interference RNAi expression was performed to validate AnxB9 and CG32032 antibodies. The appropriate UAS-RNAi lines were crossed to *wg/xg^O^-* or *10C12-*Gal4, respectively. Crosses were performed at 25°C for 2-3 days and vials were then transferred at 29°C to maximize RNAi efficiency until late L3 larvae were dissected.

### Glia isolation and annotation from scRNA-seq in the adult stage

Glial clusters were isolated from the previously published dataset (Ozel, Simon et al. 2021). We subset the clusters that were annotated as glial clusters by the authors based on high transcriptome correlation of these clusters with the bulk transcriptome of repo-positive sorted cells and low correlation with that of elav-positive sorted cells (Ozel, Simon et al. 2021). We subset the following clusters from the final annotations found in the whole optic lobe adult dataset: “G04”, “G08”, “G03”, “G09”, “G06”, “G/LQ3”, “G02”, “G05”, “G14ab”, “G/LQ1”, “G11”, “G/LQ4”, “G10”, “G07”, “G01”, “G/LQ2”, “G12”, “G13”, “G16/LQ”. Then, we analyzed subset data using Seurat package in R. Specifically, genes that exhibit variable expression in the subset were identified with FindVariableFeatures, and the data was scaled with ScaleData before principal component analysis (RunPCA). The first 20 principal components were used to embed cells in a UMAP space for visualization (RunUMAP(dims = 1:20)) and to construct a nearest neighbor graph (FindNeighbors). Re-clustering of the subset was performed with FindClusters(resolution = 0.6). This split the adult glial cells into 21 clusters, which were annotated accordingly to **Table 2**. For **Fig. S6**, we ran FindClusters(resolution = 2). (Script 2_Glia_Adult).

### Glia isolation and annotation from scRNA-seq in the L3

Glial clusters were isolated from a previously published dataset (https://github.com/NikosKonst/larva_scSeq2022/blob/main/1_preparation_of_larval_pu pal_dataset.R - larvaOL.integrated150.rds)^25^. We subset the clusters that presented a high transcriptome correlation with the bulk transcriptome of repo-positive FACS sorted cells and low correlation with that of elav-positive FACS-sorted cells^24^ (**Fig. S1A**) (Script 1b_Glia_Larva_Correlations). Then, genes that show variable expression in the subset were identified with FindVariableFeatures. Principal component analysis was performed on scaled data (ScaleData followed by RunPCA). The first 30 principal components were used to embed cells in a UMAP space (RunUMAP(dims = 1:30)) and to construct a nearest neighbor graph (FindNeighbors). The subset data was then re-clustered with a resolution of 0.8 (FindClusters(resolution = 0.8)). This split the L3 glial cells into 23 clusters, which were annotated accordingly to Table 1. (Script 1_Glia_Larva)

### Glia annotation in adult and larva

Larval glia are less studied than neurons, and we, therefore, sought specific markers to annotate and validate the clusters based on immunohistochemistry and/or RNA-FISH (**Fig. 2A**). Subperineurial glia, exclusively expressed *rost* (**Fig. 2C**), and perineurial glia expressed *svp* but not *rost* (**Fig. 2C**). Satellite glia expressed *Optix* (**Fig. 2D**) but did not express *Dll*, while lamina neuropil glia expressed both *Optix* and *Dll* (**Fig. 2D** and **E**).

- Chiasm glia (aka giant glia) expressed *CG14598* and *DAT*, a dopamine transporter responsible for the uptake of dopamine from synaptic sites (**Fig. 2F** and **6P**). However, chiasm glia can be divided anatomically into two glia subtypes: outer chiasm (xgo) and inner chiasm (xgi). We manually subset and re-analyzed the chiasm cluster in order to separate it into two unsupervised clusters. We obtained two clusters that differentially expressed *hth* and were able to identify the two clusters as being xgo (*hth*-low) and xgi (*hth*-high) (**Fig. 2G**).

- Medulla neuropil glia were originally identified in three different closely associated clusters that changed gene expression rapidly through time. This may be the consequence of these glia originating from neuroblasts after their neurogenesis window. They then migrate past all neuron cell bodies to contact the neuropil. We merged the three clusters of medulla neuropil glia into one that exhibited expression of *Hand* and *Dll* (**Fig. 2E** and **H**).

- Lobula complex neuropil glia were validated by the expression of *Cpr51A* (**Fig. 2I**) *CG4288* (**Fig. 2J**) and *Dll* (**Fig. 2E**).

- Cortex glia expressed *CG4288* (aka *MFS9*) (**Fig. 2J**), and *Cyp4g15* (**Fig. 6C**). Optic lobe cortex glia could be distinguished from central-brain cortex glia that expressed *Npc2b* (**Fig. 2L**).

- Additionally, we validated another two clusters as glial cells originating from the central brain based on their expression of *Adgf-D* (**Fig. 2M**) and their high correlation with each other (**Fig. 2B**). These central-brain glial clusters, together with the cortex central brain cluster, were excluded from the present analysis that focuses on the optic lobe.

- Wrapping glia (wg) was identified by the expression of *alphaTub85E* (**Fig. 2K**). The observed cluster corresponds to cell projections from wg located in the eye imaginal disc, as discussed above.

In the adult, we could rely more on known markers from the literature (**Fig. 3A**). For example, we could identify perineurial glia based on the expression of *Tret1-1*^46^ (**Fig. 3C**) and *Vmat*^18^, which is also expressed in fenestrated glia^49^ (**Fig. 3D**); subperineurial glia, as well as in the larva, exclusively expresses *rost* (**Fig. 3E**), while pseudocartridge glia share the same subperineurial type markers but do not express *rost*, being the subperineurial glia from the lamina. Chalice glia expressed *CG3036* and *Optix* (**Fig. 3F**). Distal satellite glia was validated by the expression of *vvl* and *b* but neither *Optix* nor *CG31663*, while proximal satellite glia expressed *vvl*, *b*, *Optix*, and *CG31663* (**Fig. 3F, G *and* J**). Cortex clusters were annotated based on the expression of wrapper^46^ and validated with *CG31663* (**Fig. 3J**). Both outer and inner chiasm expressed *CG31663*, while outer chiasm expressed *DAT* and inner chiasm expressed *GstT4* and *ome* (**Fig. 3H** and **K**). Both Epithelial and Marginal glia expresses *Optix*, these clusters were manually separated and compared to the transcriptome of FAC-sorted epithelial and marginal glia (Script 2b_Glia_Adult_Corr_Epi_Marg), then it was validated by epithelial expressing *b* and marginal expressing *GstT4* and *ome* (**Fig. 3G, H** and **K**). Neuropil glia (astrocytes-like and ensheathing) are harder to separate and annotate because it is difficult to find specific markers that are exclusively expressed in one cluster compared to the other. We found two clusters of ensheathing that we annotated as medulla ensheathing based on the expression of known ensheathing glia markers, such as *CG9657*^20^ (more markers in **Table 2**), these clusters expressed *CG31663* (also around lobula/lobula plate), *nkd*, and *ome* (both only around medulla, not lobula/lobula plate), which we found expressed in some of the medulla neuropil glia but not all. The medulla astrocytes-like cluster we annotated based on known astrocytes-like genes expression, such as *wun2*^50,51,53^, we annotated this cluster as medulla based on *nkd* expression (**Fig. 3L**) and because our integration showed that this cluster has a common trajectory with that of medulla ensheathing glia. Other neuropil glia were annotated as astrocyte-like or ensheathing based on the same known markers mentioned above, but we could not validate their identity. Because we did not find lobula/lobula plate neuropil glia in our adult dataset and because these other glial types formed a different trajectory that of medulla neuropil, we speculate these other neuropil glia are lobula/lobula plate neuropil.

### Glia integration larva, pupa and adult

We integrated our annotated larval glia and adult datasets with glial cells from P15, P30, P50 and P70 stages. We used all clusters that were considered to be glia by the authors (Özel et al., 2021): “198”, “212”, “210”, “201”, “197”, “207”, “203”, “196”, “213”, “219”, “204”, “200”, “202”, “185”, “186”, “187”, “188”, “189”, “190”, “199”, “205”, “206”, “208”, “209”, “211”, “216”, “217”. However, in the P15 stage, there was a cluster label as being of identity “206”, which was clustering by itself after integration, with no cell from other stages clustering together. Based on our analysis using the genes found in this paper as glial markers (*repo*, *CG32032*, *GstE12* and *AnxB9*), the cluster “206” in P15 is indeed not a glial cluster, we excluded this cluster from integration. After integrating all stages, we had a cluster in the center of the UMAP which was composed of cell from many different clusters and also expressed some neuronal markers (eg. *brp*, *syt1*), we excluded this cluster (Script 3_All_Glia_Integration).

We subset medulla neuropil (**Fig. 4B** and **5**) and the unknown neuropil (**Fig. S2E, F**) from the dataset with all stages. We used these clusters representing the early trajectory to look for early neuropil differentiation genes (**Fig. S4G-I**).

To look for common markers for early medulla and lamina neuropil glia, we used the integrated data and selected the clusters representing the beginning of the trajectory and used the Seurat function FindMarkers (logfc.threshold = 1, min.diff.pct = 0.4).

### Validation of glial trajectories

To validate the divergence of unique glial cell types in L3 into different adult glial types, we used known Gal4 markers to sparsely and permanently label lamina neuropil and medulla neuropil glia in L3 and observe their adult morphologies. We crossed the FLEXAMP memory cassette (Bertet, Li et al. 2014) line which carries a tub-Gal80^TS^ with R10C12-Gal4 and R54H11-Gal4. To prevent unspecific labelling, we let flies mate for 2-3 days at 18°C and kept the crosses at 18°C, and then we shifted vials to the permissive temperature (29°C) for 12-24h. After the temperature shift, white pupae were selected and returned to 18°C until adults hatched and were dissected. In the case of Hand-T2A-Gal4, the aforementioned protocol did not yield any labeled adult glia, most likely due to a weak expression of Gal4 that was insufficient to flip out the FLEXAMP cassette. In this case, we let flies mate for 2-3 days at room temperature and vials where then incubated at 29°C until adults hatched. Even in these conditions, no more than 2-3 mng glial cells were recovered per optic lobe. For *Hand-T2A*-Gal4>>memory-lacZ glial tracing we followed the same protocol as described for FLEXAMP.

### Pseudotime analysis for medulla neuropil glia

The integrated Seurat object for medulla neuropil glia consists of 3,168 single-cells from 6 developmental stages: Larva, P15, P30, P50, P70 and Adult. We used this object to make a Monocle3-based^59^ *cell_data_set* object and projected the UMAP reduction on the same coordinates as the original mng Seurat object, allowing us to perform trajectory analysis along the developmental stages used for integration. We established a pseudotime by choosing the root of the UMAP, enriched for Larva stage cells, as the starting node. We used the Monocle3 *graph_test* function to identify genes that are modulated along the pseudotime and intersected them with annotated transcription factors (TFs) in the *Drosophila* genome^79^ to find TFs that vary in expression along mng maturation. Finally, we used k-means clustering (k=3) to group TFs expressed in early, mid and late pseudotime that correspond to different developmental stages. To find genes modulated specifically along the EG and ALG branch of mng (**Fig. 3D, E**), we choose the principal graph nodes along the respective trajectories and then repeated the K-means clustering as described above.

### Identification of Cell Processes

We used two approaches based on a comparison of UMI and TFs. Approach 1 compared each cluster (or subcluster after over-clustering adult glia) with the whole dataset. To be considered processes, a cluster/ subcluster must have significantly fewer UMIs than the dataset (t-test p.adjusted < 0.005) and significantly fewer reads mapping to TFs than the dataset (t-test p.adjusted < 0.005). For approach 2, we compared each cluster/subcluster with another cluster/subcluster. To be considered processes, a cluster/subcluster must have significantly fewer UMIs (t-test p.adjusted < 0.005) than another close related cluster/subcluster (Pearson Correlation based on 2000 variable genes in the dataset > 0.9). Additionally, this cluster/ subcluster must also have fewer reads mapping to TFs (t-test p.adjusted < 0.005) than the cluster/ subcluster that it is being compared to (Script Functions).

Besides the results described in the result section, we had four clusters that showed up as being processes of other clusters by our approach 2: Inner Chiasm (processes) -> Medulla Ensheathing (cell body), Chalice -> Perineurial_CB, Epithelial and Marginal -> Cortex, and Ensheathing_2 -> Cortex glia. We considered these events false positives since, with the exception of Ensheathing_2, all these clusters were validated as having separated cell identities (Fig. S3), and cluster Ensheathing_2 had its own markers (**Table S1**). Interestingly, out of these five clusters, only inner chiasm glia was shown as processes also using approach 1.

## Data Availability

All raw and processed transcriptome data for the larval dataset are available at the GEO under accession number GSE167266. All raw and processed transcriptome data for the adult and pupa stages are at the GEO under GSE142787.

## Code Availability

All related scripts used in this manuscript are available on GitHub https://github.com/AmandaAGF/Glia_Subcell_2023.

## Acknowledgements

We thank the members of the fly community; the Bloomington stock center for flies; F. Simon for sharing the medulla origin of glial cell types; Y.-C. Chen, E.O. Mazzoni, N.C. Leite, N. Konstantinides, I. Holguera, M.N. Özel, B. Sieriebriennikov and all members of the Desplan lab for discussions and feedback on the manuscript. This work was supported by NIH R01EY017916 and R01EY13010. A.A.G.F. was partly supported by Fleur Strand Graduate Fellowship (Biology Dept. NYU), and A.A.G.F. and R.R. were partly supported by NYU’s GSAS MacCracken Program. S.C. was partly funded by EMBO Long-Term Fellowship ALTF 319-2019.

## Author contributions

A.A.G.F conceived the project. A.A.G.F., S.C. and R.R. performed and analyzed all experiments, wrote the original draft and prepared the figures. B.C. generated the UAS-sfGFPnlsPEST-2A-B2ase lines. C.D. edited the manuscript.

