## Supplementary Figures for "A Developmental Single-Cell Atlas of the *Drosophila* Visual System Glia Reveals Cell Type Diversification and Subcellular mRNA Compartmentalization"

### Figure S1

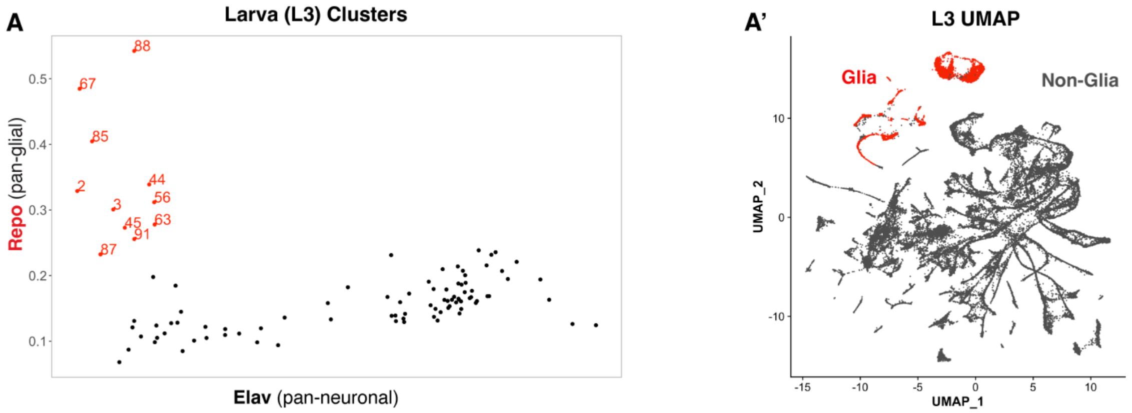


B

**Differentially Expressed Genes in Glial Compared to Non-Glial Cells**

| Gene | p_val | avg_log2FC | pct.1 | pct.2 | p_val_adj |
|---|---|---|---|---|---|
| Ama | 0 | 4.58510725 | 0.766 | 0.063 | 0 |
| MtnA | 0 | 4.26139398 | 0.675 | 0.104 | 0 |
| AnxB9 | 0 | 3.08108757 | 0.729 | 0.117 | 0 |
| CG32032 | 0 | 3.94922862 | 0.754 | 0.033 | 0 |
| Gabat | 0 | 3.07224429 | 0.698 | 0.16 | 0 |
| GstE12 | 0 | 2.59597056 | 0.625 | 0.119 | 0 |
| CG9686 | 0 | 5.08078568 | 0.642 | 0.079 | 0 |
| CG15209 | 0 | 2.91500028 | 0.556 | 0.014 | 0 |
| Npc2a | 0 | 4.38267345 | 0.752 | 0.091 | 0 |

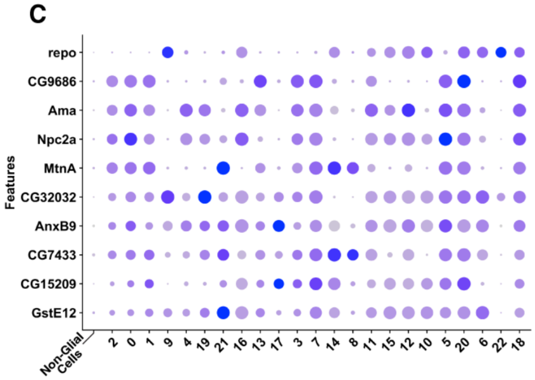


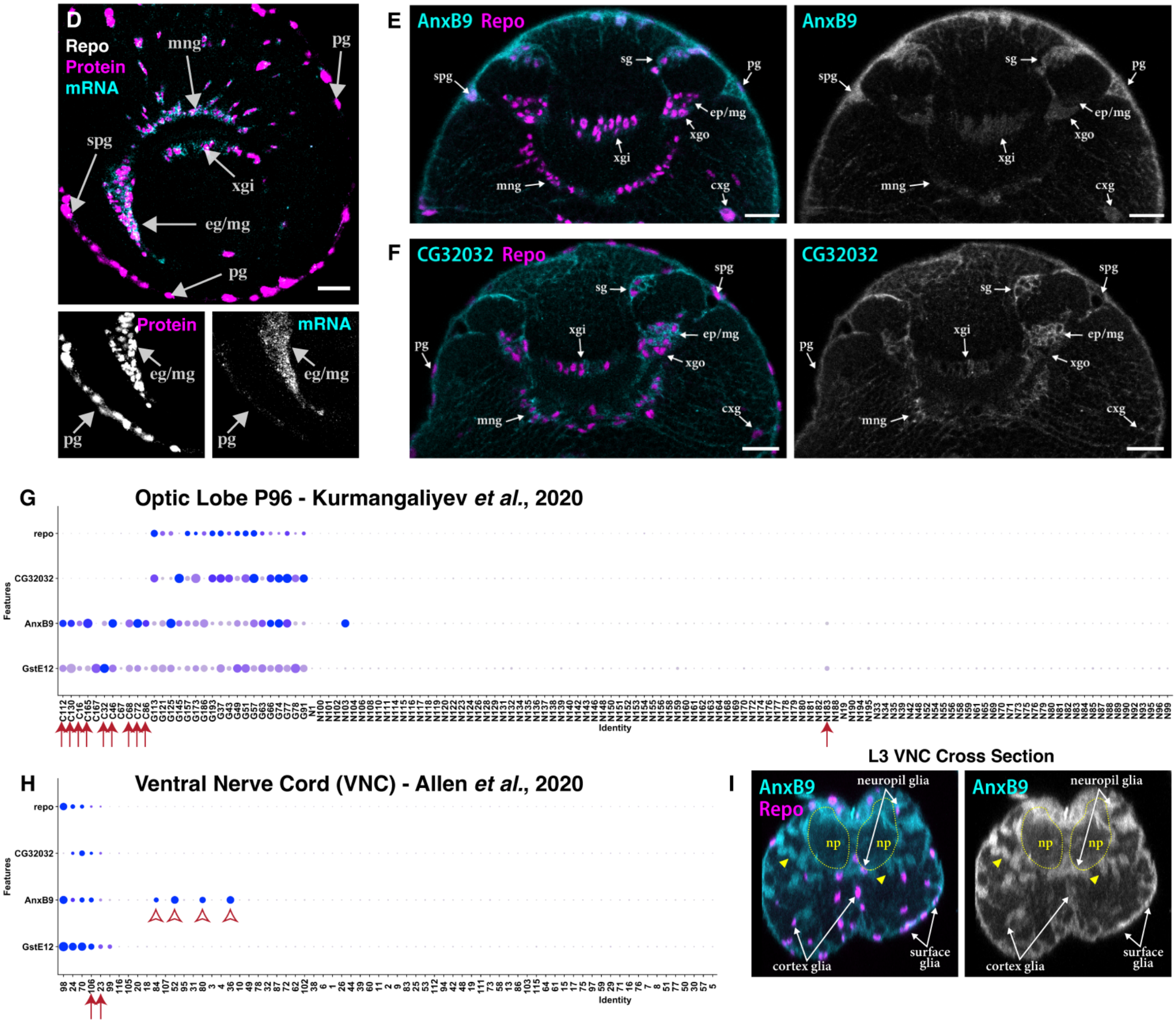

**Figure S1. Newly Identified Glia-Specific Genes.**
**(A)** Pearson correlation coefficient of larval clusters from Konstantinides *et al.*, 2022 with elav-positive bulk transcriptome (x-axis) and repo-positive transcriptome (y-axis) from Özel *et al.*, 2021. **(A')** UMAP representation of all cells in the larval dataset from Konstantinides *et al.*, 2022. Glial cells are in red and non-glial cells in dark grey.
**(B)** Table showing the top differentially expressed genes between Glial and Non-Glial cells.
**(C)** Dot plot representing the expression of all genes from **(B)** by glial clusters in the larva dataset.
**(D)** Combined HCR-RNA FISH and antibody staining for detection of both *repo* mRNA (cyan and separate channel below) and Repo protein (magenta and separate channel below). Note that surface glia presents a strong protein staining but weak mRNA signal.
**(E)** Double staining of AnxB9 (cyan and separate channel to the right) and Repo (magenta) antibodies.
**(F)** Double staining of CG32032 (cyan and separate channel to the right) and Repo (magenta) antibodies.
**(G)** Dot plot showing all non-annotated clusters from Kurmangaliyev *et al.*, 2020 – a late pupa (P96) Optic Lobe dataset. Arrows point to clusters with glial cells that were not previously identified as glia due to lack of *repo* expression.
**(H)** Dot plot showing the three newly identified glial genes by cluster of a ventral nerve cord (VNC) dataset (Allen *et al.*, 2020). Arrows point to clusters that were not identified as glia by regular pan-glial markers that were later annotated as glia by specific cell type markers by the authors. Open arrowheads show clusters that express high levels of *AnxB9* that are most likely not glia, see **(I)**.
**(I)** Cross section of a larval VNC stained for AnxB9 (cyan and separate channel to the right) and Repo (magenta). Note that some AnxB9$^{+}$ in the cortex do not present Repo antibody (yellow arrowheads) and may be neurons.
**pg** = perineurial glia, **spg** = subperineurial glia, **eg** = epithelial glia, **mg** = marginal glia, **mng** = medulla neuropil glia, **xgi** = inner chiasm glia and **np** = neuropil.

### Figure S2

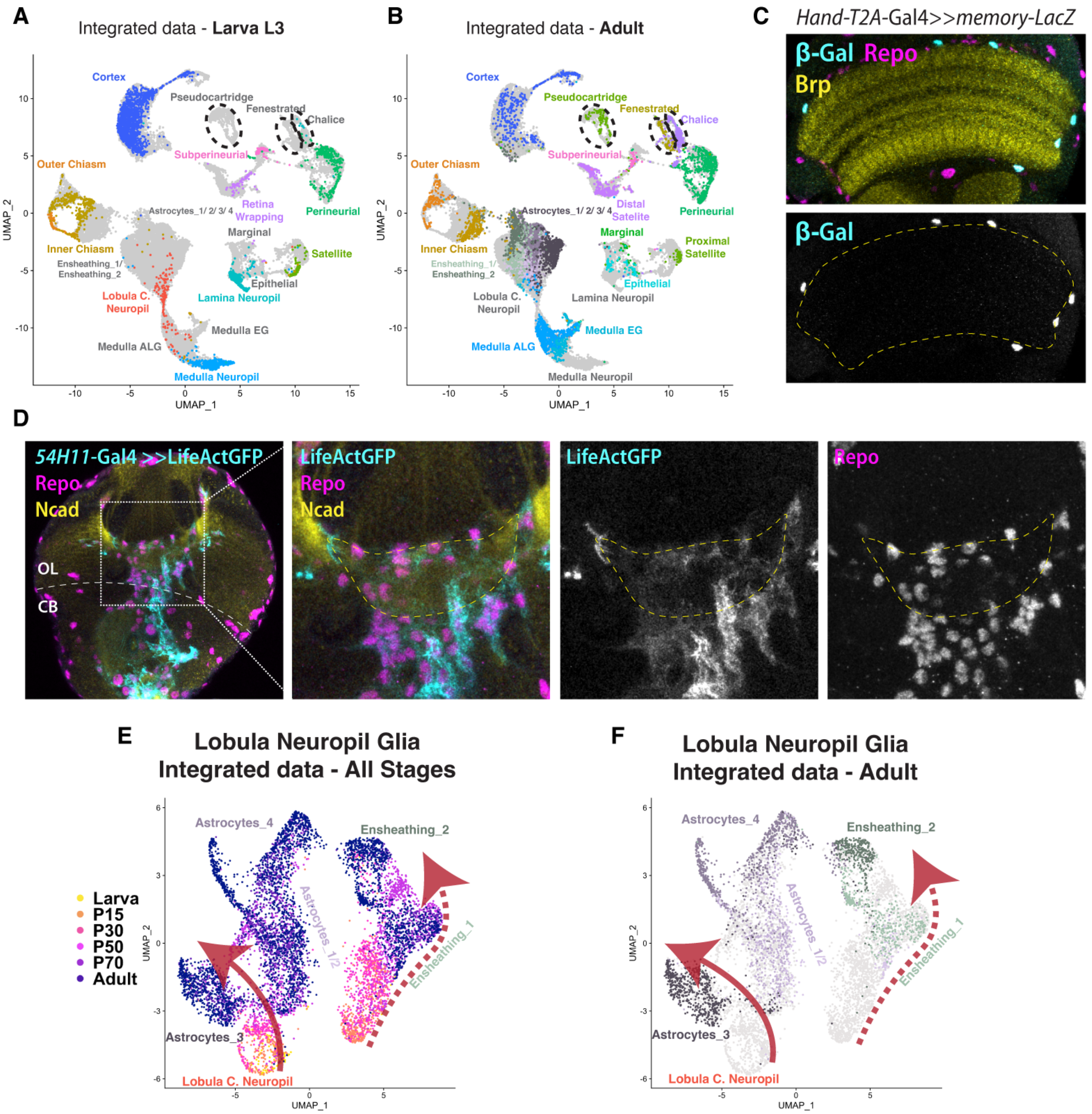
A
Integrated data - Larva L3
B
Integrated data - Adult
C
Hand-T2A-Gal4>>memory-LacZ
β-Gal Repo
Brp
β-Gal
D
54H11-Gal4 >>LifeActGFP
Repo
Ncad
OL
CB
LifeActGFP
Repo
Ncad
LifeActGFP
Repo
E
Lobula Neuropil Glia
Integrated data - All Stages
Larva
P15
P30
P50
P70
Adult
F
Lobula Neuropil Glia
Integrated data - Adult

**Figure S2. Cluster Identity in the Integrated Dataset and the Origin of Lobula Neuropil Glia.**
**(A, B)** UMAP representing the integration of all glial cells from L3, P15, P30, P50, P70 and adult stages (see **Fig. 4A**). Only larval cells are colored in **(A)**, and adult cells in **(B)**. Note that pseudocartridge, fenestrated and chalice glia are not present in the L3 dataset (dashed circles) and the expansion of neuropil glia cell types.
**(C)** *Hand-T2A*-Gal4 drives the expression of a *memory-lacZ* cassette. β-Gal (cyan and separate channel below) positive cells all express Repo (magenta), and are located in the proximal, lateral and distal surfaces of the medulla neuropil, labeled by anti-Brp antibody (yellow and yellow dashed line below).
**(D)** Section of a third instar larva optic lobe showing the lobula complex neuropil (Ncad, yellow and dashed contour). The border between the central brain (CB) and the optic lobe (OL) is shown by a dashed line. The expression of UAS-LifeActGFP (cyan, and separate channel to the right) under the control of the *54H11*-Gal4 driver labels numerous cells that cross from the CB towards the OL and associate with the lobula complex neuropil. Repo antibody (magenta, and separate channel to the right) is used to label glial cells.
**(E, F)** UMAP showing the integration of the clusters we identified as lobula neuropil glia. Cells are colored by their developmental stage of origin in **(E)** and their adult identities in **(F)**. Note that astrocyte-like clusters (Astrocytes_1/2/3/4) derive from the validated L3 lobula complex neuropil glia cluster (red arrow), whereas ensheathing glia (Ensheathing_1/2) are only represented by cells from P15 onwards (dashed-line arrow).

### Figure S3

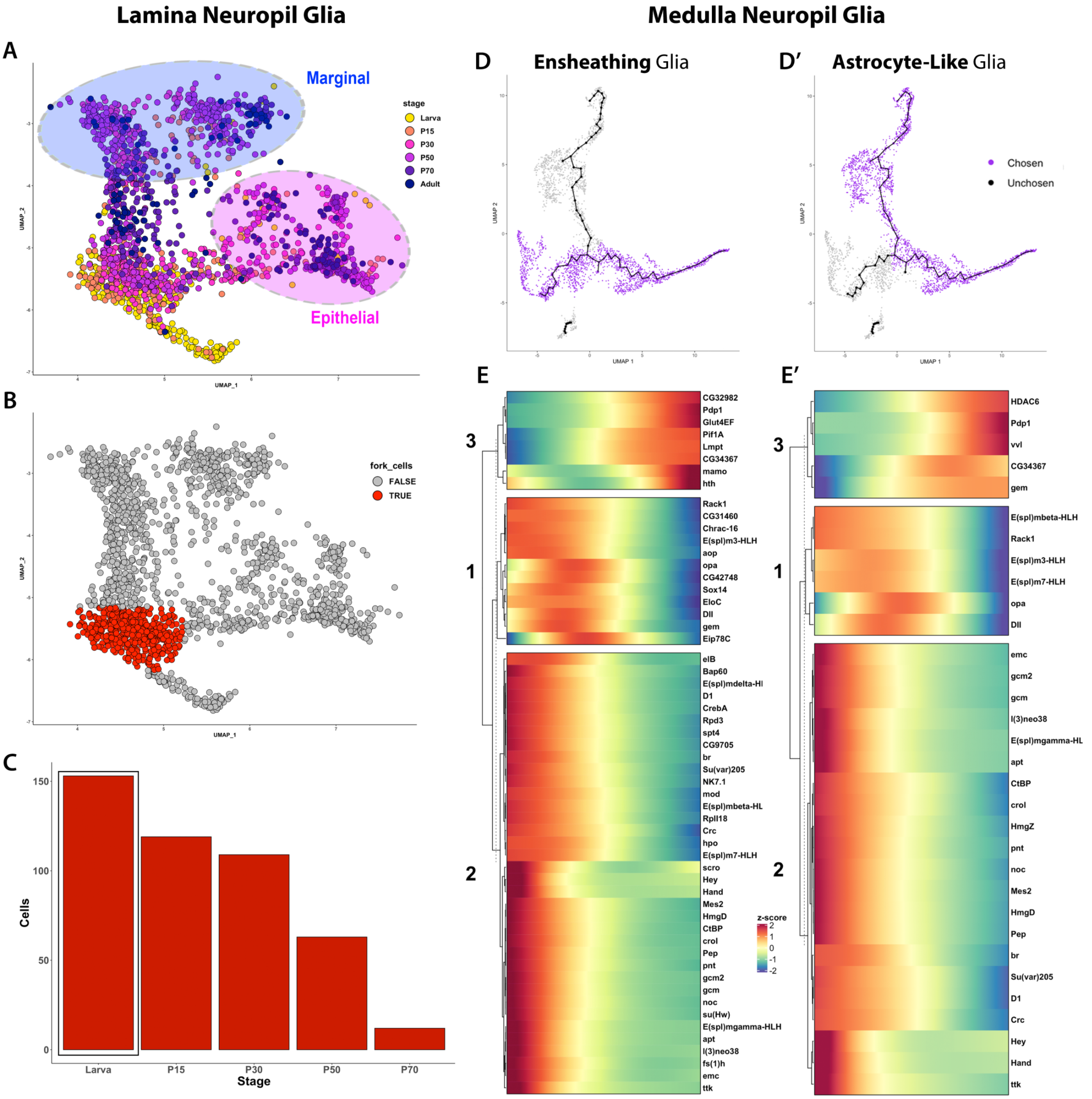
Lamina Neuropil Glia
Medulla Neuropil Glia
A
Marginal
Epithelial
stage
Larva
P15
P30
P50
P70
Adult
UMAP_1
UMAP_2
B
fork_cells
FALSE
TRUE
C
Cells
Stage
Larva
P15
P30
P50
P70
D
Ensheathing Glia
D′
Astrocyte-Like Glia
Chosen
Unchosen
UMAP 1
UMAP 2
E
CG32982
Pdp1
Glut4EF
Pif1A
Lmpt
CG34367
mamo
hth
Rack1
CG31460
Chrac-16
E(spl)m3-HLH
aop
opa
CG42748
Sox14
EloC
Dll
gem
Eip78C
elB
Bap60
E(spl)mdelta-HI
D1
CrebA
Rpd3
spt4
CG9705
br
Su(var)205
NK7.1
mod
E(spl)mbeta-HL
RpII18
Crc
hpo
E(spl)m7-HLH
scro
Hey
Hand
Mes2
HmgD
CtBP
crol
Pep
pnt
gcm2
gcm
noc
su(Hw)
E(spl)mgamma-HLH
apt
l(3)neo38
fs(1)h
emc
ttk
z-score
E′
HDAC6
Pdp1
vvl
CG34367
gem
E(spl)mbeta-HLH
Rack1
E(spl)m3-HLH
E(spl)m7-HLH
opa
Dll
emc
gcm2
gcm
l(3)neo38
E(spl)mgamma-HL
apt
CtBP
crol
HmgZ
pnt
noc
Mes2
HmgD
Pep
br
Su(var)205
D1
Crc
Hey
Hand
ttk

**Figure S3. Comparison of lamina and medulla glia lineages across development**

**(A)** UMAP embedding of isolated lamina neuropil glia integrated from all developmental stages (L3, P15, P30, P50, P70 and adult). Cells are colored by their stage of origin, showing the progressive divergence between epithelial and marginal glia fates (circled) through development.

**(B)** UMAP-based subsetting of maturing lamina neuropil glial cells at the bifurcation (colored red) of epithelial and marginal glia adult fates.

**(C)** Bar graph showing the proportion of red cells in **(B)** that belong to each developmental stage, showing that most cells at the fork belong to L3 and P15.

**(D, D')** UMAP-based trajectory selection (purple cells) illustrating maturation paths for medulla neuropil ALG and EG fates.

**(E, E')** Heatmap showing transcription factors with differential expression at early (right) mid (center) and late (left) pseudotime points for medulla neuropil EG **(E)** and medulla ALG glia **(E')** developmental trajectories.

### Figure S4

#### All Glia Clusters

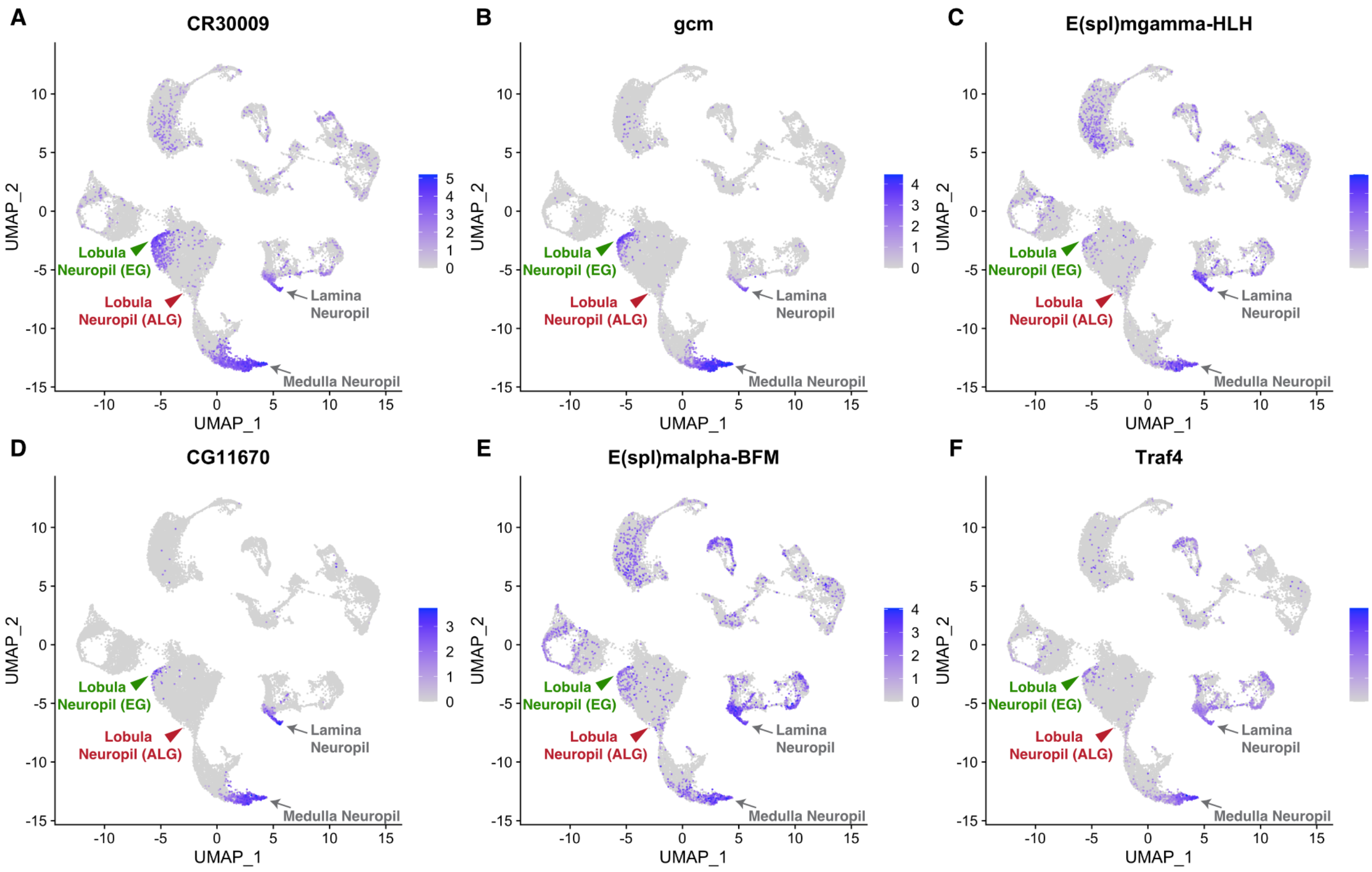


#### Lobula Neuropil Glia

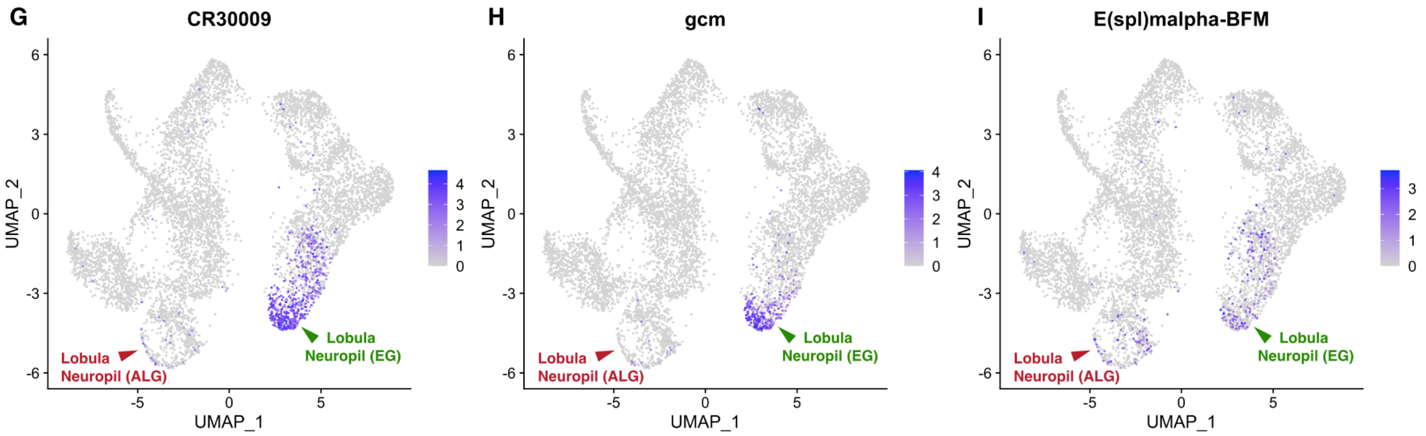

**Figure S4. Common Genes Expressed in Early Neuropil Glia.**
**(A-F)** UMAPs representing the integration of all glial cells from L3, P15, P30, P50, P70 and adult stages. Panels show the expression levels of genes that are common to the initial trajectory of neuropil glia lineages. Genes shown were the first six genes selected from **Table 3** by highest avg_log2FC and pct.2 < 0.1 (cells from other clusters that expresses the same marker). Cells expressing the genes are shown on top of the other cells (order = T).
**(G-I)** UMAPs showing only lobula neuropil glia integrated from all stages, displaying the expression levels of *CR30009*, *gcm* and *E(spl)malpha-BFM*. Cells expressing the genes are shown on top of the other cells (order = T).
The proposed start for the lobula neuropil ALG (red arrow) and lobula neuropil EG (green arrow) glia are shown in each panel. Note the relative absence of early neuropil glia markers from the lobula neuropil ALG trajectory.

### Figure S5

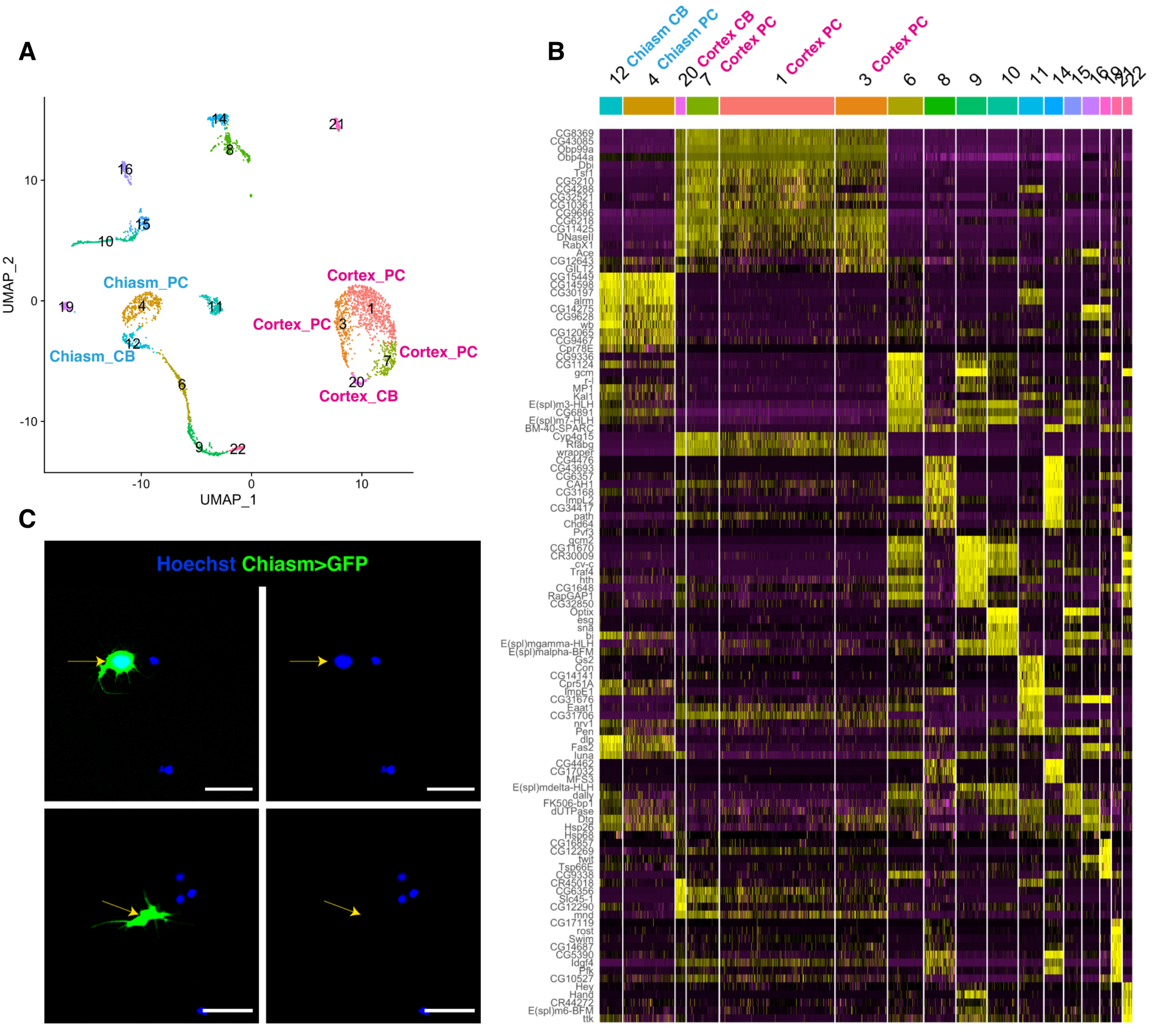

A
B
C
Chiasm_PC
Chiasm_CB
Cortex_PC
Cortex_CB
UMAP_1
UMAP_2
Hoechst Chiasm>GFP
12 Chiasm CB
4 Chiasm PC
20 Cortex CB
7 Cortex PC
1 Cortex PC
3 Cortex PC
6
8
9
10
11
14
15
16
19
21
22
CG8369
CG43085
Obp99a
Obp44a
Dbi
Tsf1
CG5210
CG4288
CG32521
CG10361
CG9686
CG6218
CG11425
DNaseII
RabX1
Ace
CG12643
GILT2
CG15449
CG14598
CG30197
alrm
CG14275
CG9628
wb
CG12065
CG9467
Cpr78E
CG9336
CG1124
gcm
r-l
MP1
Kal1
E(spl)m3-HLH
CG6891
E(spl)m7-HLH
BM-40-SPARC
Cyp4g15
Rfabg
wrapper
CG4476
CG43693
CG6357
CAH1
CG3168
ImpL2
CG34417
path
Chd64
Pvf3
gcm2
CG11670
CR30009
cv-c
Traf4
hth
CG1648
RapGAP1
CG32850
Optix
esg
sna
bi
E(spl)mgamma-HLH
E(spl)malpha-BFM
Gs2
Con
CG14141
Cpr51A
ImpE1
CG31676
Eaat1
CG31706
nrv1
Pen
dlp
Fas2
luna
CG4462
CG17032
MFS3
E(spl)mdelta-HLH
dally
FK506-bp1
dUTPase
Dtg
Hsp26
Hsp68
CG16857
CG12269
twit
Tsp66E
CG9338
CR45018
CG6356
Slc45-1
CG12290
mnd
CG17119
rost
Swim
CG14687
CG5390
Idgf4
Pfk
CG10527
Hey
Hand
CR44272
E(spl)m6-BFM
ttk

**Figure S5. scRNA-seq Reveals Subcellular Distribution of mRNAs**

**(A)** UMAP representing the L3 glial clusters annotated accordingly to cluster numbers prior to annotation (**Fig. 2B** and **Table 1**). Chiasm glia clusters are highlighted in blue are cortex glia clusters in pink. PC = Cellular Processes, CB = Cell Body.

**(B)** Heatmap representing the top markers for each cluster (high expression in yellow, and low expression in purple). The chiasm glia cluster 12 and 4 are highly similar, sharing the same markers, as is the case for the cortex glia clusters 20, 7, 1 and 3.

**(C, D)** Dissociated cells from L3 optic lobe expressing membrane-bound GFP in chiasm glia. Hoechst labels cellular nuclei. Chiasm glia with nucleus **(C)** and Chiasm glia fragments without nucleus **(D)**. Scale bar = 20 mm.

### Figure S6

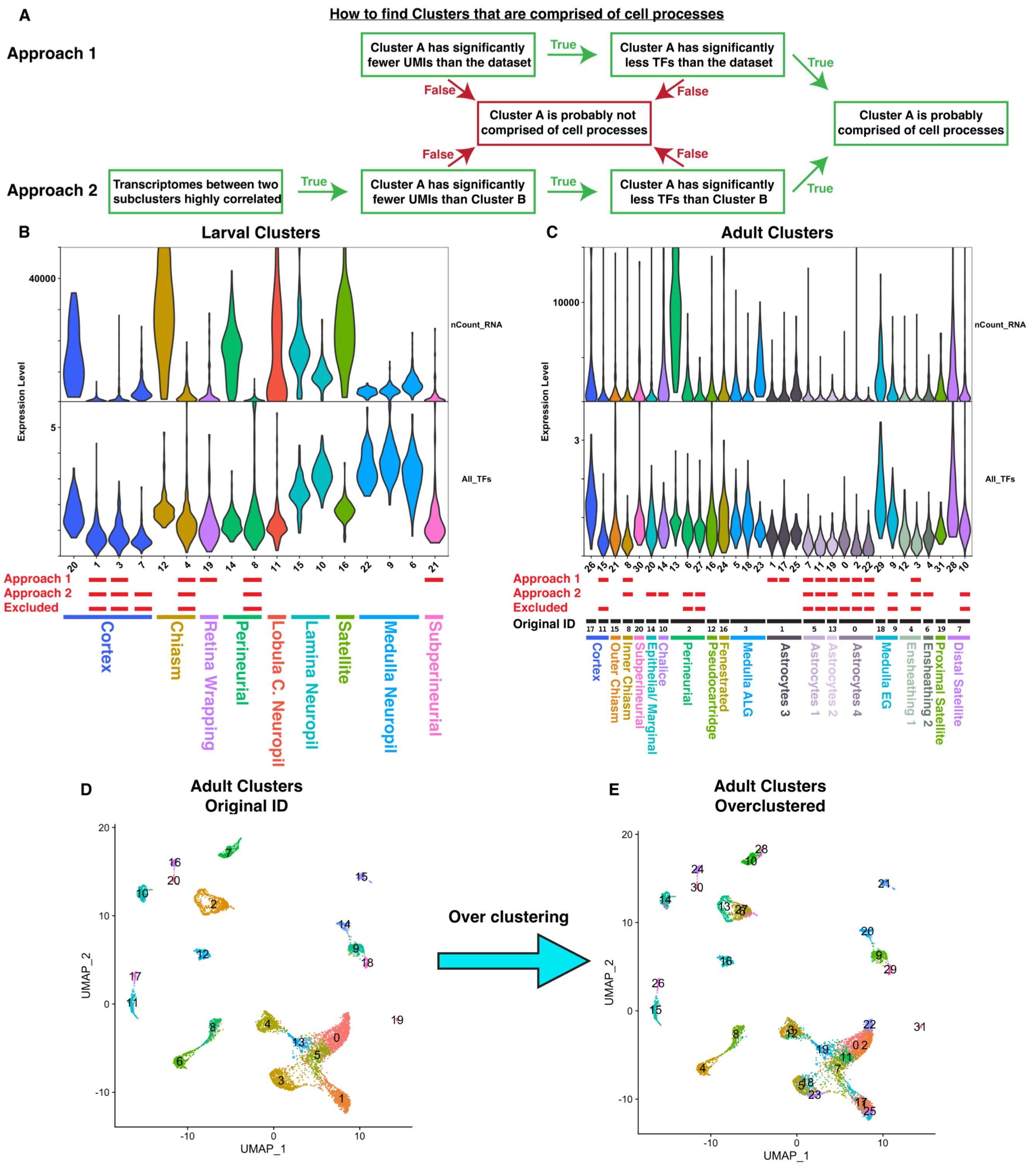
A
How to find Clusters that are comprised of cell processes
Approach 1
Cluster A has significantly fewer UMIs than the dataset
True
Cluster A has significantly less TFs than the dataset
True
False
False
Cluster A is probably not comprised of cell processes
Cluster A is probably comprised of cell processes
False
False
Approach 2
Transcriptomes between two subclusters highly correlated
True
Cluster A has significantly fewer UMIs than Cluster B
True
Cluster A has significantly less TFs than Cluster B
True
B
Larval Clusters
40000
5
Expression Level
nCount_RNA
All_TFs
20 1 3 7 12 4 19 14 8 11 15 10 16 22 9 6 21
Approach 1
Approach 2
Excluded
Cortex
Chiasm
Retina Wrapping
Perineurial
Lobula C. Neuropil
Lamina Neuropil
Satellite
Medulla Neuropil
Subperineurial
C
Adult Clusters
10000
3
Expression Level
nCount_RNA
All_TFs
26 15 21 8 30 20 14 13 6 27 16 24 5 18 23 1 17 25 7 11 19 0 2 22 29 9 12 3 4 31 28 10
Approach 1
Approach 2
Excluded
Original ID
17 11 15 8 20 14 10 2 12 16 3 1 5 13 0 18 9 4 6 19 7
Cortex
Outer Chiasm
Inner Chiasm
Subperineurial
Epithelial/ Marginal
Chalice
Perineurial
Pseudocartridge
Fenestrated
Medulla ALG
Astrocytes 3
Astrocytes 1
Astrocytes 2
Astrocytes 4
Medulla EG
Ensheathing 1
Ensheathing 2
Proximal Satellite
Distal Satellite
D
Adult Clusters
Original ID
UMAP_2
UMAP_1
Over clustering
E
Adult Clusters
Overclustered
UMAP_2
UMAP_1

**Figure S6. Computational Approach to Identify Clusters of Cellular Processes**
**(A)** Flowchart describing the two different approaches used to identify clusters comprised of cell processes. Both approaches were used in parallel in our work.
**(B)** Violin Plot showing each cluster from the larval dataset. Red bars below indicate whether each cluster was identified as processes by Approach 1, Approach 2, and if the cluster was excluded from the final atlas.
**(C)** Violin Plot showing each subcluster from the adult dataset after over-clustering. Red bars below indicate whether each subcluster was identified as processes by Approach 1, Approach 2 and if the subcluster was excluded from the final atlas. It also shows which original ID cluster the subclusters come from.
**(D, E)** UMAP plots showing the original ID **(D)** and the over-clustering **(E)** of the adult dataset.

### Figure S7

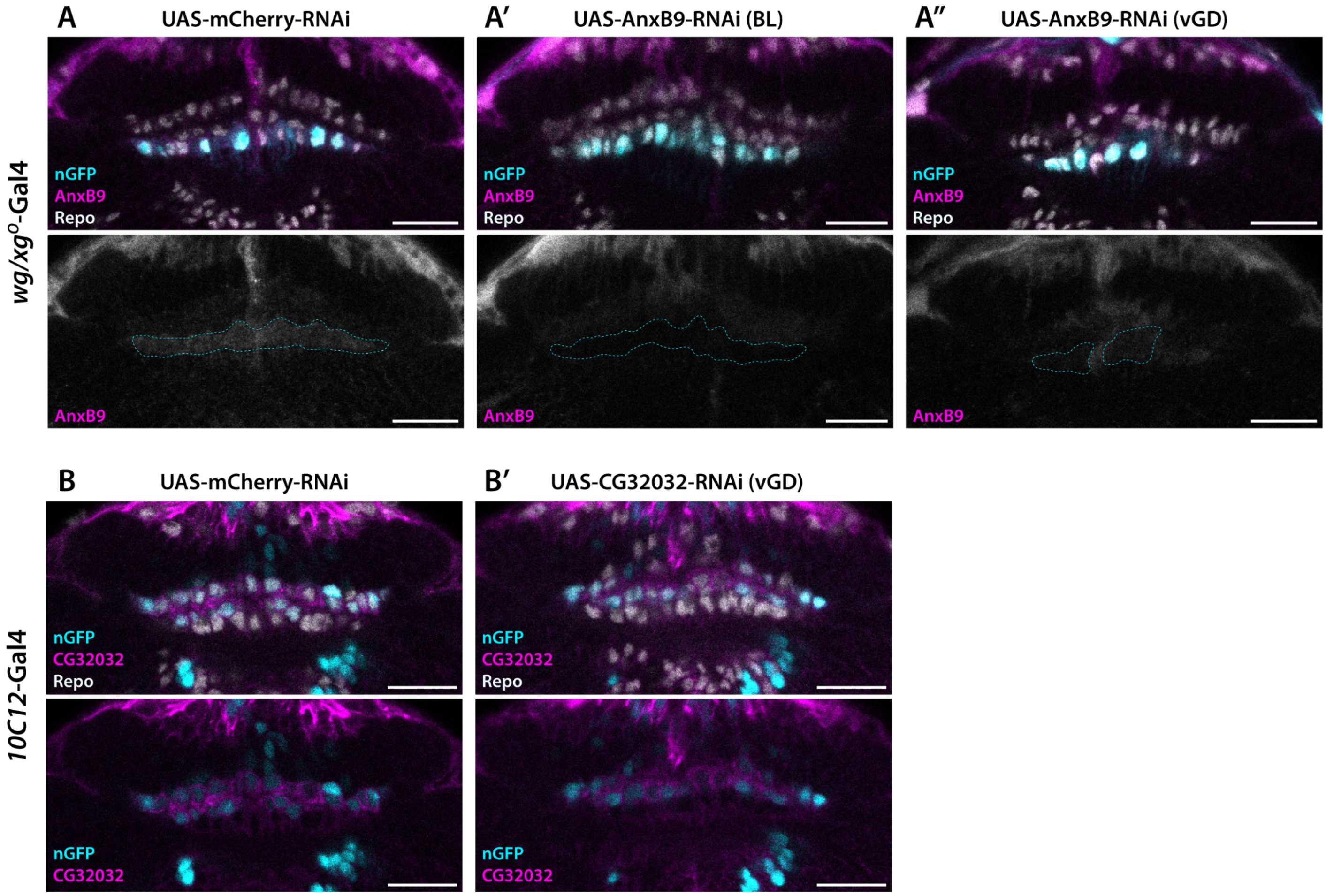
A
UAS-mCherry-RNAi
A′
UAS-AnxB9-RNAi (BL)
A″
UAS-AnxB9-RNAi (vGD)
wg/xg^O-Gal4
nGFP
AnxB9
Repo
AnxB9
B
UAS-mCherry-RNAi
B′
UAS-CG32032-RNAi (vGD)
10C12-Gal4
nGFP
CG32032
Repo
nGFP
CG32032

**Figure S7. Validation of AnxB9 and CG32032 Antibodies.**
**(A)** *wg/xg$^{O}$*-Gal4 driver is used to express UAS-nlsGFP-PEST (cyan) in wrapping and outer chiasma glia (cyan dashed contour in lower panels), which show higher levels of AnxB9 antibody (magenta and separate channel below). Repo (grey) is used to identify glial cells. Control UAS-mCherry-RNAi **(A)**, and two different interference RNA constructs against *AnxB9* **(A', A'')** are expressed, showing a cell type-specific reduction of AnxB9 signal in outer chiasma glia.
**(B)** *10C12*-Gal4 driver is used to express UAS-nlsGFP-PEST (cyan) in lamina neuropil glia, which display highe levels of CG32032 antibody. We did not observe any difference in CG32032 signal in optic lobes expressing UAS-mCherry-RNAi controls **(B)** or a UAS-CG32032-RNAi construct **(B')** in lamina neuropil glia.
